# Binding Free Energies of Piezo1 Channel Agonists at Protein-Membrane Interface

**DOI:** 10.1101/2022.06.27.497657

**Authors:** Wenjuan Jiang, Han Zhang, Yichun-Lin, Wonpil Im, Jerome J. Lacroix, Yun Lyna Luo

**Affiliations:** Department of Pharmaceutical Sciences, Western University of Health Sciences, California, US; Department of Basic Medical Sciences, Western University of Health Sciences, California, US; Department of Biological Sciences, Chemistry, Bioengineering, and Computer Science and Engineering, Lehigh University, Pennsylvania, US

## Abstract

Mechanosensitive Piezo channels convert mechanical stimuli into biological signals in vertebrates. Piezo1 chemical modulators are anticipated to yield many clinical benefits. To date, Yoda1 is the most potent and widely used Piezo1-selective agonist, yet how Yoda1 interacts with Piezo1 at the protein-membrane interface and stabilizes Piezo1’s open state remains elusive. Here, using a previously identified putative Yoda1 binding site and three molecular dynamics (MD)-based methods, we computed the binding free energies of Yoda1 and its analogs in a Piezo1 cryo-EM closed state and an *in silico* open state. Our computed absolute binding free energy of Yoda1 in the closed state agrees well with the experimental *K*_d_ in which Piezo1 is expected to be in a closed state. More importantly, Yoda1 binds the open state better than the closed state, in agreement with its agonist effects. All three methods predicted that Dooku1, a Yoda1 analog, binds the closed state stronger than Yoda1, but binds the open state weaker than Yoda1. These results are consistent with the fact that Dooku1 antagonizes the effects of Yoda1 but lacks the ability to activate Piezo1. The relative binding free energies of seven Yoda1 analogs recapitulate key experimental structure-activity-relationships (SAR). Based on the state-dependent binding free energies, we were able to predict whether a molecule is an agonist or inhibitor and whether a chemical modification will lead to a change in affinity or efficacy. These mechanistic insights and computational workflow designed for transmembrane binders open an avenue to structural-based screening and design of novel Piezo1 agonists and inhibitors.

## Introduction

Developing drugs that act as highly ion channel-specific modulators remains challenging. Small molecules targeting ion channels may diffuse to the pore from the solution (hydrophilic pathway) or partition into membrane before entering binding sites (lipophilic pathway). Through the lipophilic pathway, drugs may access to channel pores through membrane-exposed fenestration or bind between transmembrane (TM) helices at the protein-membrane interface^1^. Decades of drug design efforts have focused on the binding at the protein-aqueous phase, but much less attention has been given to ligands binding at the protein-membrane interface (we call them TM binders). This delay is largely due to the technical challenges in identifying and quantifying TM binders. In addition, the intrinsic hydrophobicity of TM binders may cause promiscuous binding if the target protein consists of large TM domains. In other words, there may exist multiple silent binding sites in the TM region with no functional outcome.

Those challenges in studying TM binders were manifested in the mechanosensitive Piezo ion channels. The two isoforms in vertebrates, Piezo1 and Piezo2 channels, are among the largest known plasma membrane proteins (>7600 amino acids). Their homotrimer structures harbor a central pore (TM37-38), an extracellular cap, an intracellular CTD domain, and three large TM domains (TM1-36) called blades or arms^2-6^. Each arm consists of nine transmembrane bundles of four α-helices, called Piezo Repeat A-I (or THU9-THU1)^3^. An intracellular beam lays below Repat A-C (**Fig. 1a**). In total, a single Piezo channel consists of 114 TM helices. The first Piezo1-selective small-molecule agonist named Yoda1 (C_13_H_8_Cl_2_N_4_S_2_) (**Fig. 1b**) was discovered by functional screening of 3.25 million small molecules before the Piezo1 structure was solved^7^. This seemingly large number represents only a tiny fraction of the estimated small molecule chemical space (e.g. ∼10^60^ for molecules harboring 30 heavy atoms)^8^. Thus, the knowledge of Yoda1’s binding site and reliable prediction of agonist activity on the Piezo1 channel will provide a new avenue to expand currently limited Piezo channel modulators.

**Figure 1.**
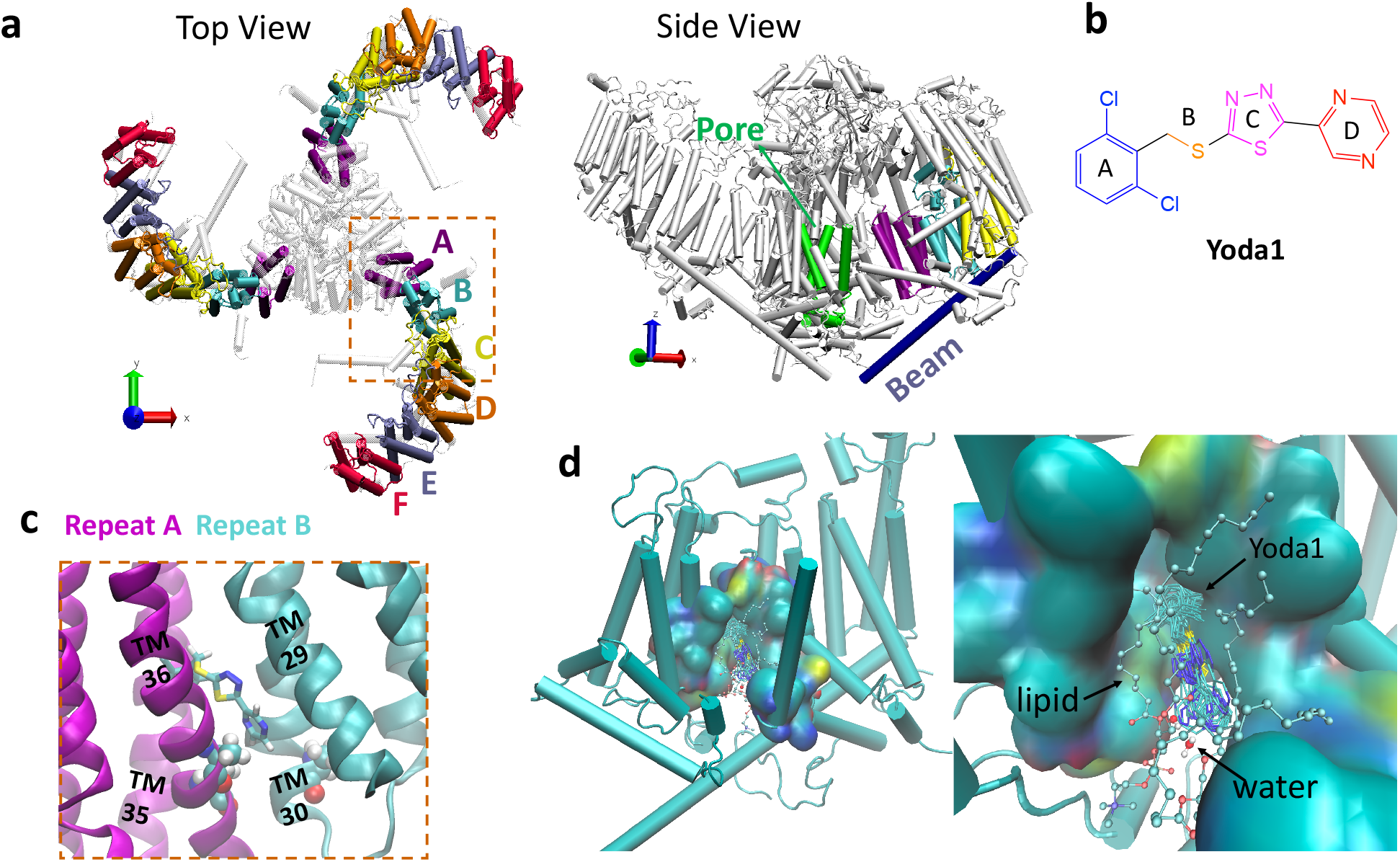
a) Structure of Piezo1 (PDBID:6B3R) and repeat A to F for each arm shown with top and side views. Pore and Beam regions are colored in green and dark blue, respectively. b) Piezo1’s agonist Yoda1. c) Yoda1 binding pose between repeat A (purple) and repeat B (cyan) in Piezo1 closed state. Transmembrane helices TM29-30 and TM35-36 are labeled. A2094 on TM36 and A1718 on TM30 are shown in vdW mode. d) Overlaid Yoda1 binding pose in the open state. Binding pocket is shown in surface mode. Lipid and water molecules within 3 Å of Yoda1 are shown in CPK mode in VMD. Atom color: cyan carbon, blue nitrogen, red oxygen, yellow sulfur.

Since its discovery, Yoda1 has been used widely as a chemical probe and greatly deepened our understanding of the biological functions of Piezo1 channels^9-21^. Clinically, pharmacological activation of Piezo1 in the vascular endothelium could mimic some of the beneficial cardiovascular effects of exercise^22^, whereas activation of Piezo1 in red blood cells could prevent malaria-causing *Plasmodium* infection^23-24^. Piezo1 activation could also help treat lymphedema^17, 20^, promote muscle regeneration^15^ and bone formation^25^. Owing to its potential therapeutic usage, chemical space around the Yoda1 scaffold has been explored by synthesis prior to the knowledge of the binding site^26-27^. It was shown that Yoda1’s agonist activity is susceptible to single atom modifications, but the underlying reason is unknown.

Electrophysiological investigations using pressure-clamp recordings reveal Yoda1 displaces Piezo1’s pressure-dependent activation curve toward lower pressure values^7^ and slows down the kinetics of Piezo1 macroscopic ionic currents during inactivation (up to ∼9-fold) and deactivation (up to ∼20-fold)^28^. Thus, Yoda1 acts as a “gating modifier” by energetically stabilizing a channel open state (conducting state) vs. closed state (nonconducting state). Yoda1-mediated Piezo1 activation is maintained when purified channels are reconstituted into droplet bilayers^7^, in absence of cellular environment. In addition, a direct interaction between Yoda1 and the purified mouse Piezo1 arm in detergent (residues 1-2190, consisting of TM1-36 helices) has been measured using surface plasma resonance (SPR) technique^29^. By engineering chimeras between mouse Piezo1 and its Yoda1-insensitive paralog Piezo2, the Repeat A region of Piezo1 (residues 1961-2063, consists of TM33-35 helices) was shown essential to mediate the agonist effects of Yoda1^19^. Based on these accumulated experimental evidence, the first all-atom molecular dynamics (MD) simulation of Piezo1 was conducted to search the Yoda1 binding site^30^. Ligand mobility analysis of 20 Yoda1 molecules over 4.8 *μs* MD trajectories revealed that Yoda1 partitions into the lipid bilayer and remains stable at the TM region sandwiched between Repeat A and B for at least 3.5 *μs* (**Fig. 1c**). Steric mutations in this binding site abrogate Yoda1-mediated Piezo1 activation *in vitro*^30^, suggesting these mutations may prevent Yoda1 from wedging in between these two Repeats. Moreover, this TM region is 40 Å distant from the Piezo1 pore, suggesting Yoda1 may act as an allosteric agonist.

Despite these collective efforts, direct structure-function evidence supporting this Yoda1 binding site as a functionally relevant agonist binding site is still lacking (i.e., a binding site ≠ an agonist binding site). Due to the large size of Piezo1 (∼8000 amino acids) and the low solubility of Yoda1, solving the structure of a Yoda1-Piezo1 complex at high resolution remains challenging. In addition, silent binding sites (no agonist effect) may exist due to the sheer size of Piezo1 (1.2 million Daltons) and the hydrophobic nature of Yoda1 (355 Daltons, *c*LogP 3.5). The Piezo structures captured by cryo-EM are commonly thought to correspond to a closed/non-conducting channel conformation populated in absence of mechanical stimulus. The curved Piezo arms are anticipated to flatten in the presence of a mechanical stimulus, promoting channel opening^29, 31-35^. Based on this hypothesis, Jiang et al. generated an open conformation of Piezo1 by imposing a membrane-protein curvature mismatch during all-atom MD simulations^33^. The gating motions of the Piezo1 arm, cap, and pore observed from simulations were beautifully recaptured by a recent cryo-EM study of the curved and flattened structures of Pieoz1 reconstituted in small liposomes^36^. The open pore recapitulates experimentally obtained ionic selectivity, unitary conductance, and mutant phenotypes^33^. This Piezo1 open state thus provided a new opportunity for interrogating the structural and thermodynamic basis of Yoda1 agonism on Piezo1 channel.

Here, using three MD-based binding free energy calculations, conducted with three MD engines and two sets of force fields, we show that Yoda1 binds to Piezo1 open state stronger than to the closed state, consistent with its agonist activity on Piezo1. In contrast, a non-agonist Yoda1 analog, Dooku1, showed the opposite binding preference. Our state-dependent binding simulations of seven Yoda1 analogs revealed two reasons behind the loss of agonist activity by chemical modifications: affinity loss and efficacy loss. These mechanistic insights and the prediction power of the computational approaches will be useful for expanding the chemical space of Piezo1 channel modulators.

## Results and Discussion

According to the classical agonist mechanism, Yoda1 may activate Piezo1 through conformational selection and/or induced fit. Thermodynamically, the efficacy of Yoda1 on Piezo1 is equivalent to the relative binding affinity of Yoda1 in the open state vs. closed state of the Piezo1 arm (**Fig. 2)**. An agonist will have a stronger binding affinity in the open state vs. closed state 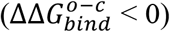, while an antagonist will bind to the closed state stronger 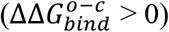. A silent binder will bind to both states undifferentiable 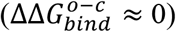. Thus, we can compare the binding free energy of a ligand in open and closed states to confirm its agonist effect without computing the free energy of Piezo1’s open to closed transition with ligand bound (*holo*) or unbound (*apo*). The latter is much more computationally expensive and technically challenging. As a comparison, we also computed the binding free energy of Dooku1, a Yoda1 analog that has no agonist effect on Piezo1, but antagonized Yoda1’s effect on Piezo1.

**Figure 2.**
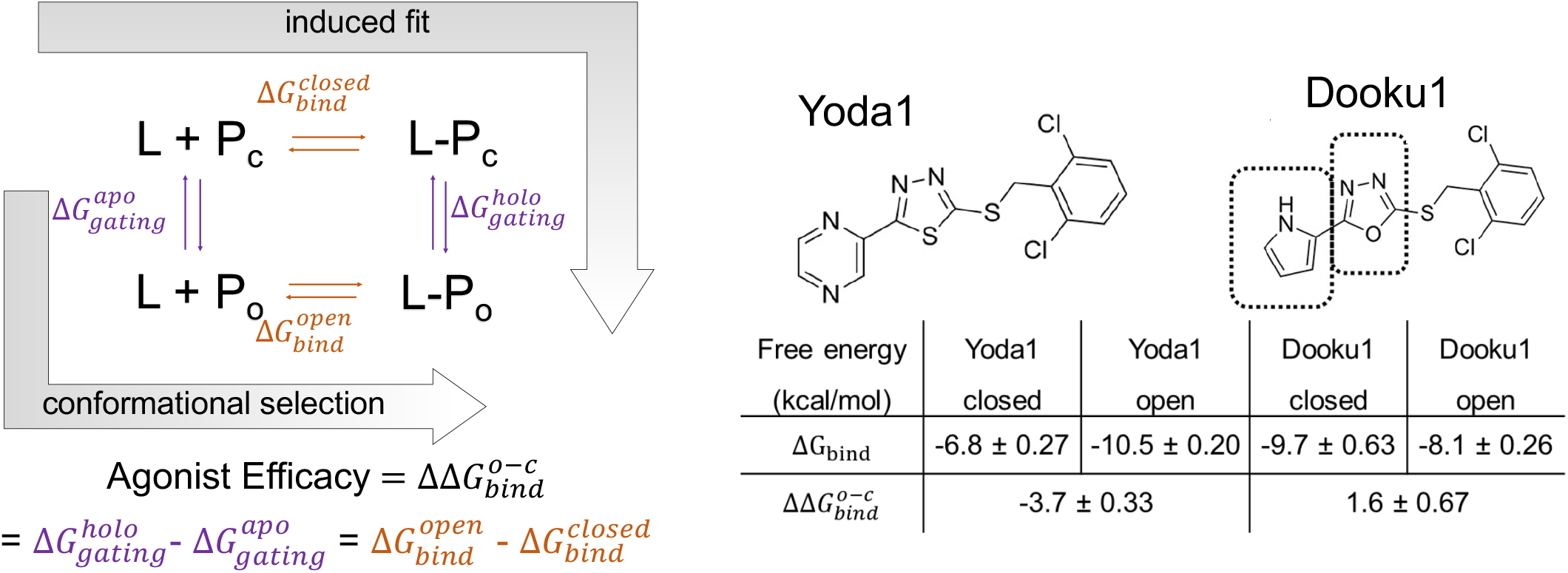
Thermodynamic framework of Piezo1 agonist Yoda1 and non-agonist Dooku1. **a)**. Thermodynamic cycle links ligand (**L**) efficacy to relative binding affinities in the two functional states of the protein (**P**). **P**_**o**_ represents the open (active) state and **P**_**c**_ represents the closed (deactivated or inactivated) state of the protein. **b)** Absolute binding free energy (ABFE) calculations predict an agonist role for Yoda1 but not for its non-active analog Dooku1.

Three computational approaches (summarized in **Table 2**) were used in this study to compare the binding affinities of Yoda1 and Dooku1 in Piezo1 open and closed states: 1) the absolute (standard) binding free energy (ABFE),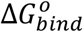, refers to the Gibbs free energy difference at binding site vs. in bulk solution in 1M standard concentration; 2) relative binding free energies (RBFE) of Yoda1 vs. analogs; 3) Site Identification by Ligand Competitive Saturation (SILCS). Those simulations were carried out using three MD simulation engines (NAMD, AMBER,GROMACS), and two sets of force fields for proteins (CHARMM36 force field, AMBER ff19SB) and small molecules (CGenFF with dihedral fitting, GAFF2.1), respectively.

### Unbiased simulations

Five replicas of 100 ns of unbiased MD simulations were carried out for four systems, namely Yoda1-open, Dooku1-open, Yoda1-closed, and Dooku1-closed (open and closed indicate the conformation of Piezo1). During all simulations, Yoda1 and Dooku’s common dichlorophenyl moiety (ring A) is more stable at the hydrophobic binding site and in contact with 1-2 lipid tails from the inner leaflet (**Fig. 1d**). In contrast, the pyrazine moiety in Yoda1 or pyrrole in Dooku1 (ring D) is mobile and forms dynamical contacts with lower leaflet lipid headgroups and water. These observations indicate that 1) dichlorophenyl is critical for binding; 2) the mobile pyrazine side of the Yoda1 scaffold should tolerate more chemical modification towards better efficacy and water solubility. This relative “loose” protein-ligand complex may represent a unique feature of hydrophobic (“greasy”) molecules binding at the TM region, in sharp contrast to the ligands anchored by hydrogen bonds at the protein-water interface.

### Absolute binding free energy (ABFE)

ABFE is equivalent to the reversible thermodynamic work needed to move a ligand from its binding pocket to the bulk solution. It allows a direct comparison with the experimental binding affinity measured based on ligand concentration in solution. Since free energy is a state function, alchemical free energy calculations can be used to compute ABFE via scaling the ligand-environment interactions using a thermodynamic coupling parameter *λ* from 0 to 1. Alchemical approaches often rely on predefined geometric restraints between protein and ligand to avoid the so-called “wandering ligand” problem when interactions are scaled down. In theory, ABFE does not depend on the details of the restraints if the free energy contribution of each restraint is accounted for. In practice, if the restraints prevent a complete sampling of the ligand at the end states, it can give results that are highly reproducible but incorrect. Hence, to ensure the correct sampling of the “greasy” TM-binders, we used the sampling information from 5×100 ns unbiased MD to define the upper boundaries of two half flat-bottom harmonic restraints to ensure unbiased sampling of fully coupled state and efficient sampling in the uncoupled state (see **Theory and Methods**, and **Fig. 5**). The FEP coupled with replica-exchange MD (FEP/REMD) implemented in NAMD^37^ was used for computing all ABFEs, in which nonbonded interactions between the ligand and its environment are scaled at the binding site and in bulk solution (i.e. double-decoupling scheme)^38^. Comparing the sampling at the fully coupled stage (*λ* = 1) with unbiased MD (**Fig. 5**), we saw that ligands are less likely to be kinetically trapped during FEP/REMD thanks to the frequent exchange between *λ* stages (see Convergence of ABFE section).

The final ABFE results in **Figure 2** (detailed in **Table S1)** indicate that Yoda1 binds to Piezo1 open state -3.7 ± 0.3 kcal/mol stronger than to the closed state, consistent with electrophysiological data showing that Yoda1 stabilizes Piezo1 open state, hence acting as an agonist. ABFE results further show that Dooku1 binds to the Piezo1 closed state stronger than Yoda1, but its binding affinity in the open state is weaker than Yoda1. Thus, our state-dependent ABFE results are entirely consistent with the experimental observation that Dooku1 inhibits Yoda1’s agonist activity but lacks agonist activity on Piezo1 channel. In fact, our data suggests Dooku1 stabilizes the Piezo1 closed state by 1.6 ± 0.67 kcal/mol, to a much lesser degree than Yoda1’s stabilization on the Piezo1 open state. Although it is currently not clear whether there exists an experimental assay sensitive enough to capture Dooku1’s inhibitory activity on Piezo1, our results opened an opportunity for designing Piezo1 inhibitors that favor Piezo1 closed state over open state.

In Piezo1 open state, Yoda1 binds ∼3 Å lower (towards the intracellular side) than Dooku1 in the same binding region (**Fig. 5**). To further rule out the possibility that Dooku1 could bind stronger if it reaches the deeper pocket as Yoda1, we manually switched Yoda1 and Dooku1 binding positions and conducted two additional ABFE calculations for which we call Yoda1-open-switch and Dooku1-open-switch (**Fig. S7**). The ABFE results (**Table S1**) show that the original lower binding site is more favorable for Yoda1, while Dooku1 binds the upper and lower site similarly. Overall, Yoda1 binds the open state stronger than Dooku1 even in their switched binding poses.

Experimentally, Yoda1’s effective concentration EC_50_ from in cellular to in vivo assays are ∼10-50 *μ*M^7, 39^. The binding affinity *K*_d_ between Yoda1 and the purified mouse Piezo1 from SPR binding assay is 45.6 ± 14.3 *μ*M^29^, roughly corresponding to -6.0 kcal/mol at room temperature. This value agrees well with our calculated ABFE of -6.8 ± 0.2 kcal/mol in Piezo1 closed system, consistent with the fact that purified Piezo1 in detergent is likely a closed or similar to closed conformation.

### Relative binding free energy (RBFE)

Our state-dependent ABFE calculations not only reproduced experimental *K*_d_ of Yoda1, but also provided quantitative mechanistic insights behind the functional outcomes of Yoda1 and Dooku1. Yet, these calculations have low throughput. ABFE calculations require decoupling the whole ligand from its environment and need to be repeated for each ligand and in both open and closed states. For small modifications of a known chemical scaffold, RBFE calculations are more efficient. We hence tested whether RBFE can capture the difference in agonist activity between Yoda1 and its analogs. Using Yoda1 as the reference compound, we define RBFE as 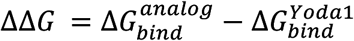 Thus, analogs having ΔΔ*G<* 0 are predicted to bind stronger than Yoda1, and vice versa. In addition, ΔΔΔ*G*^*o*−*c*^ (= ΔΔ*G*^*open*^ *−* ΔΔ*G*^*closed*^) corresponds to the efficacy of the analog relative to Yoda1. As expected from a thermodynamic loop (**Fig. 2**), ligands having ΔΔΔ*G*^*o*−*c*^ *<* 0 are predicted to be better agonists than Yoda1, while ligands having large positive ΔΔΔ*G*^*o-c*^ can potentially act as Piezo1 inhibitors.

We tested this idea using seven known Yoda1 analogs from literatures whose agonist activities on mouse Piezo1 are available^26, 40^. RBFE simulations were conducted using GPU accelerated AMBER-TI method with AMBER ff19SB force field (protein) and GAFF2.1 (ligand). RBFE results indicate Dooku1 binds to the Piezo1 open state weaker than Yoda1 (ΔΔ*G*^*open*^ > 0) but binds to the closed state stronger than Yoda1 (ΔΔ*G*^*closed*^ *<* 0) (**Table 1**), similar to ABFE predictions (**Fig. 2**). This agreement between RBFE and ABFE, i.e. two independent methods and force fields, strongly supports our computational approach. Comparing analogs **7b** and **11** it is evident that both pyrrole rings and the oxadiazole ring in Dooku1 reduce binding affinity relative to Yoda1 in the open state, but pyrrole ring alone makes Dooku1 and analog **7b** binds to closed state stronger. Replacing ring D with phenyl surprisingly renders the analog **2i** binds stronger in both states; however, the efficacy is lost. Oxidation of thioether to sulfone (**1b**) is unfavorable in both states. The trajectory of **1b** show that the bulky sulfone prevents the tight binding of dichlorophenyl ring. Removing two chlorine atoms on ring A reduces binding affinity (**1a**), while adding a fluorine at meta-position slightly increase the binding affinity in the open state (**2a**). These RBFE predicted SARs are in excellent agreement with the qualitative experimental observations^26, 40^.

**Table 1.**
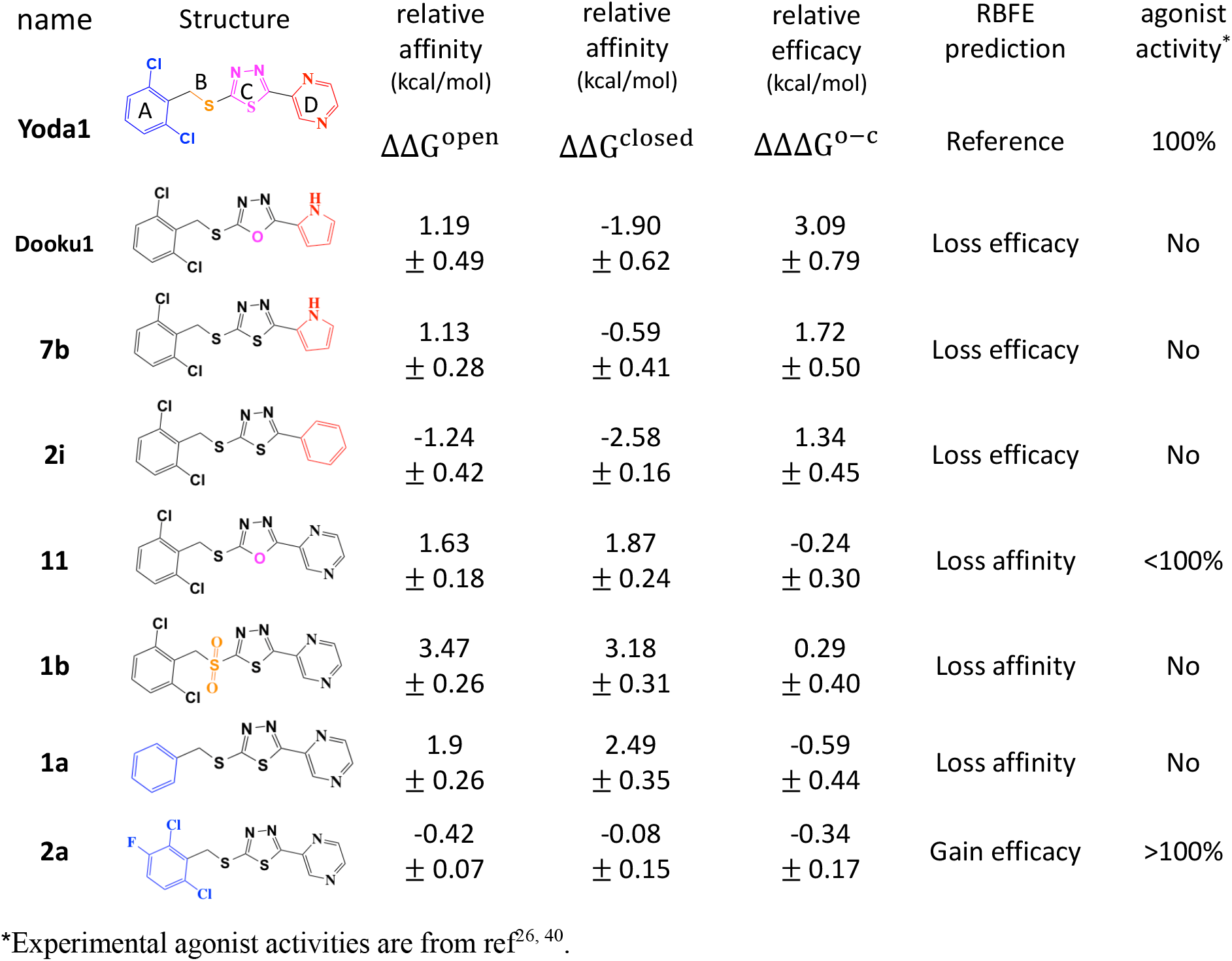
Relative binding free energy results of Yoda1 analogs. Standard deviations are computed from four replicas of 5-10 ns AMBER-TI runs.

| name | Structure | relative<br>affinity<br>(kcal/mol) | relative<br>affinity<br>(kcal/mol) | relative<br>efficacy<br>(kcal/mol) | RBFE<br>prediction | agonist<br>activity* |
| --- | --- | --- | --- | --- | --- | --- |
| Yoda1 | | $\Delta\Delta G^{open}$ | $\Delta\Delta G^{closed}$ | $\Delta\Delta\Delta G^{o-c}$ | Reference | 100% |
| Dooku1 | | 1.19<br>$\pm 0.49$ | -1.90<br>$\pm 0.62$ | 3.09<br>$\pm 0.79$ | Loss efficacy | No |
| 7b | | 1.13<br>$\pm 0.28$ | -0.59<br>$\pm 0.41$ | 1.72<br>$\pm 0.50$ | Loss efficacy | No |
| 2i | | -1.24<br>$\pm 0.42$ | -2.58<br>$\pm 0.16$ | 1.34<br>$\pm 0.45$ | Loss efficacy | No |
| 11 | | 1.63<br>$\pm 0.18$ | 1.87<br>$\pm 0.24$ | -0.24<br>$\pm 0.30$ | Loss affinity | <100% |
| 1b | | 3.47<br>$\pm 0.26$ | 3.18<br>$\pm 0.31$ | 0.29<br>$\pm 0.40$ | Loss affinity | No |
| 1a | | 1.9<br>$\pm 0.26$ | 2.49<br>$\pm 0.35$ | -0.59<br>$\pm 0.44$ | Loss affinity | No |
| 2a | | -0.42<br>$\pm 0.07$ | -0.08<br>$\pm 0.15$ | -0.34<br>$\pm 0.17$ | Gain efficacy | >100% |

\*Experimental agonist activities are from ref<sup>26, 40</sup>.

### Site Identification by Ligand Competitive Saturation (SILCS)

In addition to ABFE and RBFE approaches, we seek to validate Yoda1 binding site using an orthogonal approach that does not require prior knowledge of the binding site. We have learnt from our ABFE simulations that lipids are absent in the binding region in Piezo1 closed state, but present in open state. In contrast, there are more water molecules in the closed state binding site in proximity to the lipid headgroup region than in the open state (**Fig. S8**). Thus, the dynamics of protein, lipids, and water molecules at the Yoda1 binding region is critical for accurate prediction of binding affinity. It is hence not surprising that conventional docking scoring functions benchmarked on soluble proteins has led to very poor predictions when tested on ion channels^41^. SILCS method in heart is a set of saturated binding simulations of small drug fragments. It uses grand canonical Monte Carlo/MD (GCMC/MD) method that oscillates chemical potential to accelerate the fragment binding/unbinding events, thus can generate complex fragment density maps (FragMaps) of target proteins in a realistic membrane environment^42^. The newly added halogen FragMaps feature made SILCS an appealing approach for testing Yoda1 binding site.

The FragMaps of Piezo1 arm were generated from ten copies of 100 ns SILCS GCMC/MD simulations. Ligand docking to FragMaps is performed using the SILCS-Monte Carlo (SILCS-MC) sampling method, which consists of energy minimization, MC moves (translation, rotation, dihedral rotation), and MC simulated annealing to obtain the lowest ligand grid free energy (LGFE) binding pose. Five independent SILCS-MC samplings were conducted, and the results are considered converged when the lowest LGFE scores are within 0.5 kcal/mol. Using SILCS-MC global docking, we found that the lowest LGFE pose of Yoda1 matches perfectly with our ABFE simulation. At this position, Yoda1’s dichlorophenyl matches the density of chlorobenzene and fluorobenzene (**Fig. 3**). Thus, it is not surprising that removing chlorine substituents reduces binding affinity and adding fluorine increases affinity. Most importantly, LGFE score also predicted that Yoda1 binds stronger in open state (−9.8 kcal/mol) than in closed state (−8.8 kcal/mol), while Dooku1 binds closed state slightly stronger than open state (−9.4 vs. -9.2 kcal/mol).

**Figure 3.**
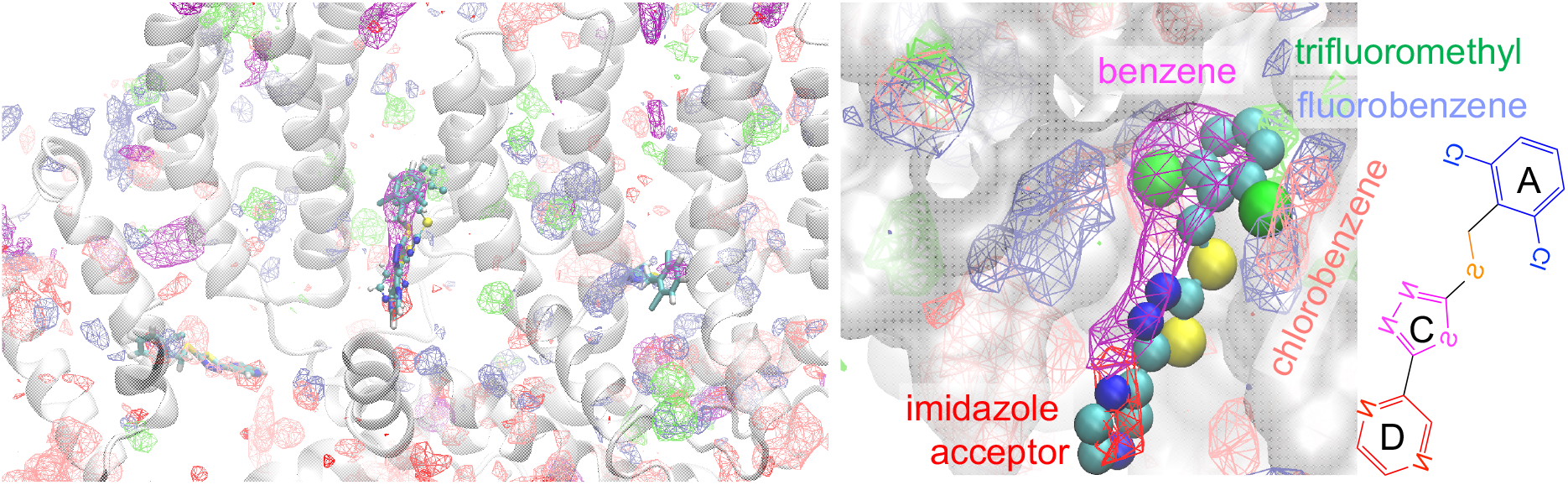
SILCS FragMaps of Piezo1 open state and the top binding pose of Yoda1 generated from SILCS-MC docking.

### Conclusion and Perspectives

Drugs that positively or negatively modulate Piezo1 channel functions are highly sought after. However, the sheer size of Piezo1 channel and the lack of structural information of a potential drug binding site have hindered the rational drug design for this important target. Our previous MD simulations of a cryo-EM closed state have identified a putative Yoda1 binding site at Piezo1’s TM region between Piezo1 repeat A and B. Steric mutations Ala*→*Trp in this binding pocket abrogate the effects of Yoda1. The location of this pocket is consistent with our prior study showing that replacing part of Piezo1 repeat A by its homologous counterpart from Piezo2 abolishes the effects of Yoda1^19^. However, direct structural evidence of this binding site is still lacking. Our *in silico* Piezo1 open state model gives us the opportunity to interrogate the thermodynamic framework of Yoda1.

Using three different computational binding free energy calculations and two different force fields (**Table 2**), our data consistently show that Yoda1 binds stronger in Piezo1 open state, while its analog Dooku1, favors Piezo1 closed state. This mechanistic insight unraveled the possibility of this TM region as agonists and inhibitors binding site. During all simulations, Yoda1 is interacting with lipid and water molecules at the TM binding site. Besides the dichlorophenyl moiety, the rest of the Yoda1 molecule is highly mobile. To capture this dynamic binding feature at TM region, we implemented a set of flat-bottom restraints to allow unbiased sampling of fully coupled ligands during FEP/REMD. Our ABFE results demonstrated that structural stability is not necessary for thermodynamic stability. The binding affinity computed in the closed state is in remarkable agreement with the *K*_d_ measured experimentally, consistent with the expectation that Piezo1 is in closed state when measured in detergent. Our analysis suggests that flat-bottom restraints are suitable for computing ABFE of highly mobile ligands and can help optimize the binding pose when a ligand bound crystal structure is unavailable.

**Table 2.**
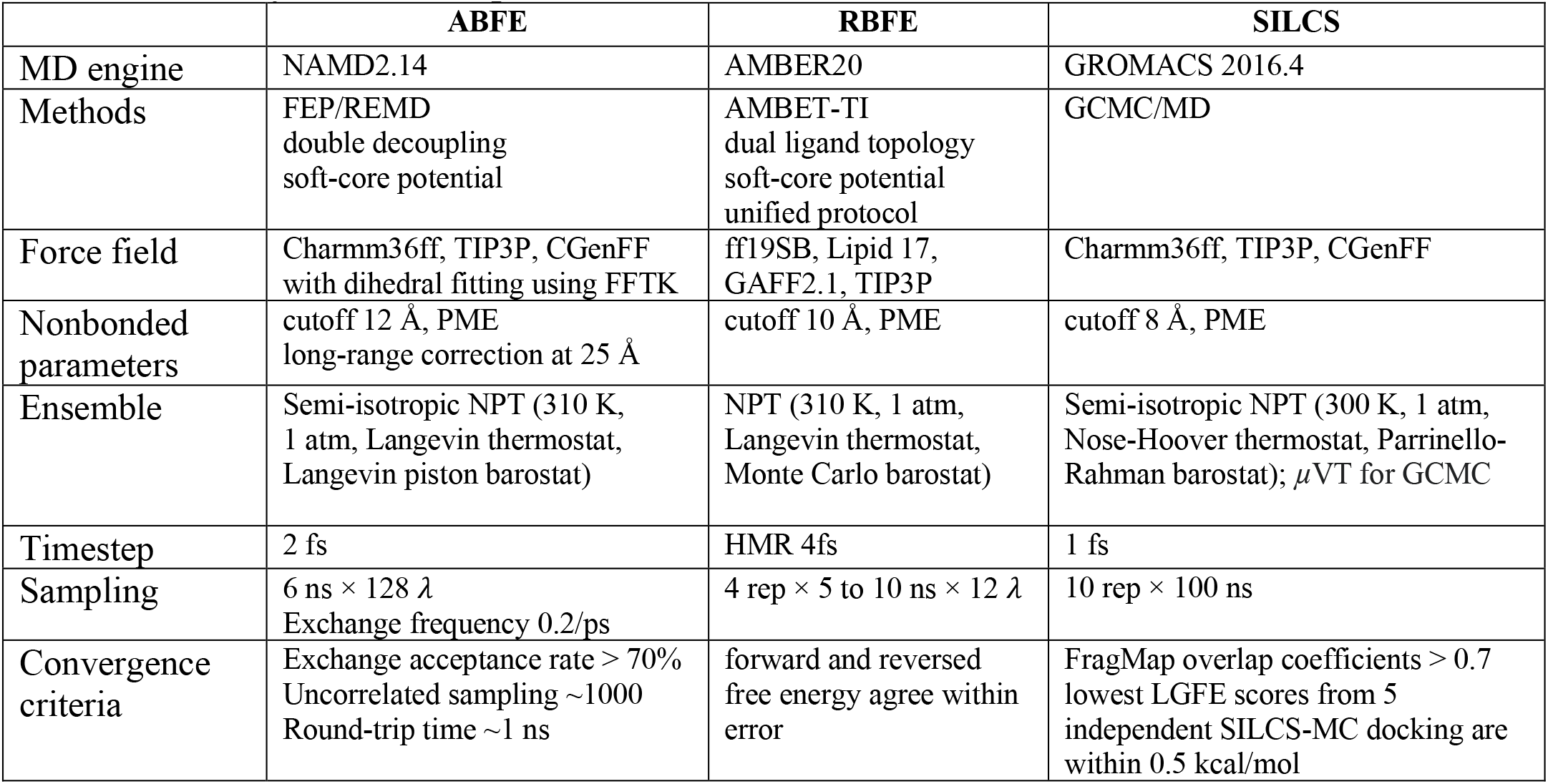
Summary of three computational methods.

The SAR of Yoda1 analogs was further interrogated using the GPU accelerated AMBER-TI. The RBFE of each ligand in Piezo1 open and closed conformation provided the mechanistic insight behind the loss or gain of the agonist activity. Unlike ABFE and RBFE simulations, SILCS method does not require prior knowledge of binding site. Based on the FragMaps of Piezo1 in open and closed states, SILCS-MC docking reproduced faithfully the Yoda1 binding pose in our ABFE and RBFE simulations. This agreement is likely because SILCS FragMaps are generated in presence of lipid environment. The mechanistic insights and the prediction power of the computational approaches presented here will be useful for expanding the chemical space of Piezo1 channel modulators.

Interestingly, we also observed that although the three computational methods produced qualitative agreement that Yoda1 binds to the open state stronger than Dooku1 and Dooku1 binds to the closed state stronger than Yoda1, the sensitivity of the three methods in this case is decreasing as the computational methods becomes less expensive (**Fig. 4**). Therefore, SILCS-MC docking can serve as initial screening to search new compound scaffolds. As more compound data become available, SILCS has the potential to be combined with machine learning algorithms to screen billion compound libraries for ion channels. In addition, the atomic level of details offered by visual analysis of the FragMaps provides invaluable insights that can facilitate our decision making in a medicinal chemistry campaign.

**Figure 4.**
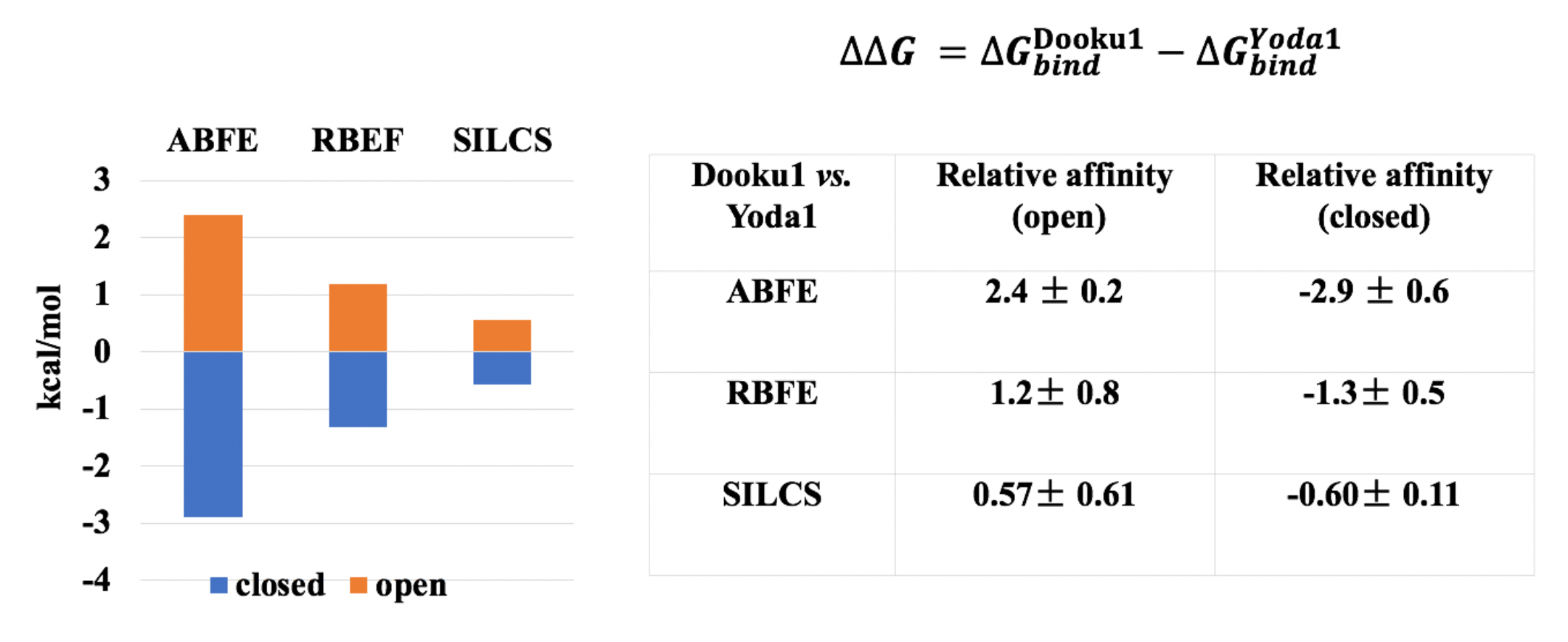
Comparison of the relative binding affinity of Dooku1 vs. Yoda1 in Piezo1 open and closed states from three computational methods.

## Theory and Methods

Three MD-based computational methods were used to compare the binding affinities of Yoda1 and its analogs in Piezo1 open and closed states. **Table2** provides a brief summary of each method. Please see each section for more details.

### Absolute binding free energy (ABFE) simulations

#### System setup

Yoda1 binding pose in closed Piezo1 conformation was taken from the last frame of previous 4.8 *μs* MD trajectory^30^. The initial binding pose of Yoda1 in Piezo1 open conformation was generated by aligning the four TM helices (TM29-30 on Repeat B, and TM35-36 on Repeat A, residue 1683 to 1733 and 2047 to 2097) consisting of the binding site. The Piezo1 open state was taken the last frame of 2 µs trajectory^33^. Dooku1 binding pose is generated by modifying Yoda1 structure in the binding site using MOE software^43^. Since the binding affinity is influenced by local protein conformation and local membrane/solvent environment, a single Piezo1 arm (residue 1131 to 2190) embedded in the lipid bilayer is used instead of the whole Piezo1 trimer (**Fig. 1**). To keep the monomer stable, weak backbone RMSD restraints were applied on the first and last helices of the arm remote from Yoda1 binding site. Each protein-ligand complexes were prepared in solvated POPC bilayer using CHARMM-GUI^44-45^. CHARMM36 parameter sets were used for the protein^46^, ions^47^, lipids^48^, and TIP3P for water^49^. CHARMM general force field (CGenFF)^50-51^ was used for Yoda1 and Dooku1 molecules, with dihedral fitting using Force Field Toolkit plugin^52^ of the VMD 1.9.3^53^. The force fields and input files are available on Github (https://github.com/reneejiang/ABFE-inputs). The terminal amino acids of Piezo1 segments are capped using acetylated N-terminus (ACE) and methyl-amidated C-terminus (CT3) blocking groups. The standard MD equilibrium and production simulation protocols are detailed in **Supplementary Methods**. Five independent replicas of 100 ns unbiased MD were first conducted using AMBER20 pmemd.cuda on RTX2080Ti GPU cards for each ligand-protein complex.

#### Flat-bottom harmonic restraint setup

Due to the high mobility of Yoda1 binding at transmembrane region, we used two flat-bottom harmonic restraints during FEP/REMD to ensure unbiased sampling of fully coupled state and efficient sampling in uncoupled state. To do so, two ensemble distributions were computed from a total of 500 ns unbiased trajectories: 1) the distance *R* between center of mass (COM) of ligand and the binding site, and 2) ligand conformational *RMSD* after rigid-body alignment of the binding site to the initial snapshot, i.e., distance-to-bound-configuration (DBC) in NAMD colvars^54^. The boundaries of the flat-bottom harmonic restraint for ligand conformation *U*_*c*_ and protein-ligand distance *U*_*d*_ (force constant 100 kcal/mol/Å^2^) are set to be beyond the limit of the ensemble distribution of *R* and *RMSD* so that when the ligand is fully coupled (*λ*=1), the free energy contribution of the *R* and *RMSD* restraints are negligible (**Fig. 5**).

**Figure 5.**
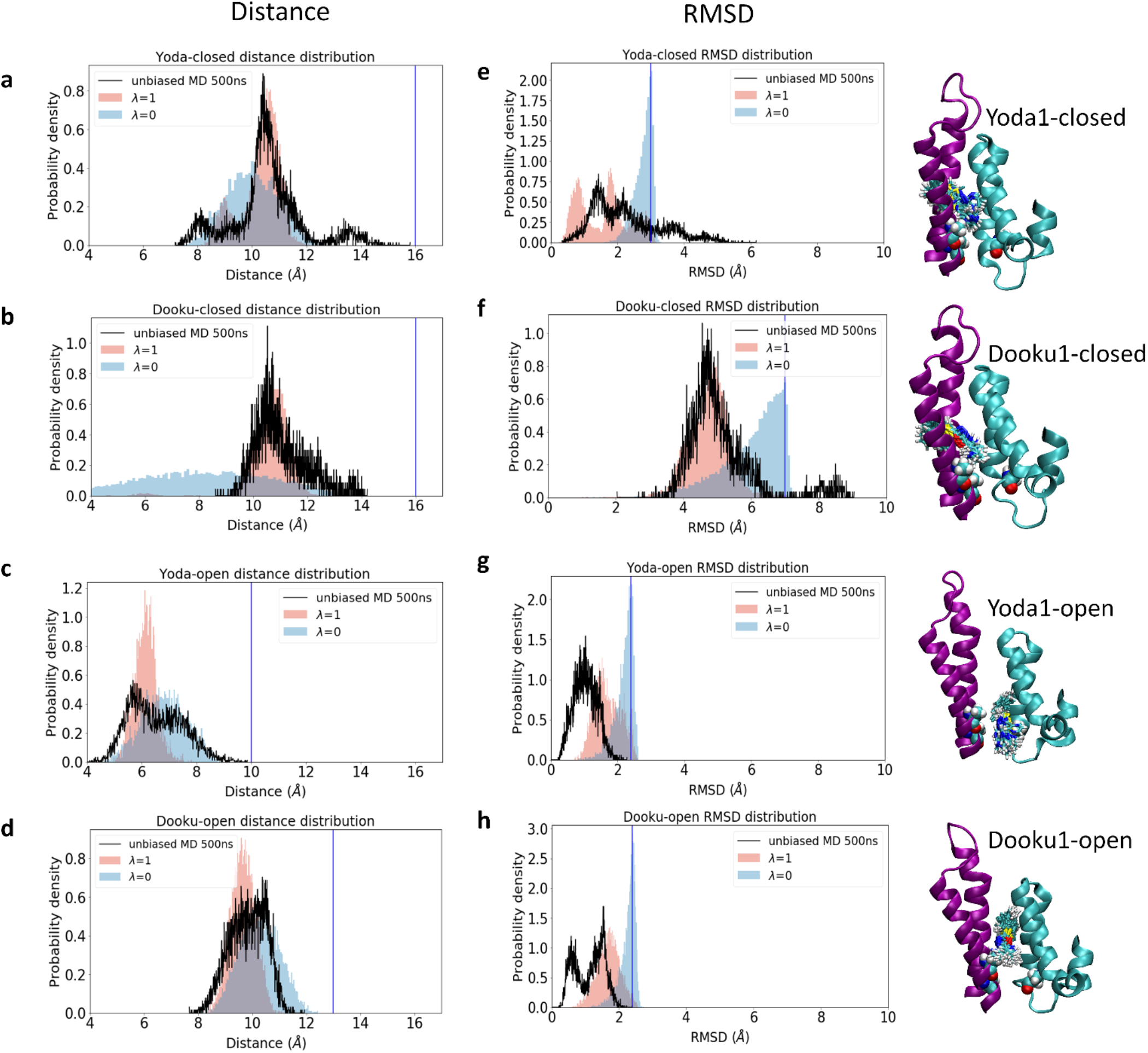
**abcd.** The distributions of the protein-ligand distance *R* for Yoda1 and Dooku1 in Piezo1 closed and open states. **efgh**. The distributions of ligand conformational *RMSD* after rigid-body alignment of the binding site to the initial snapshot, for Yoda1 and Dooku1 in Piezo1 closed and open states. In each plot, distributions from unbiased simulations are shown in black; distributions from fully coupled (*λ* =1) FEP/REMD trajectories are shown in pink; distribution from fully uncoupled (*λ* =0) trajectories are shown in blue. Blue vertical lines indicate the boundaries of the half flat-bottom harmonic restraint for ligand conformation *U*_*c*_ and protein-ligand distance *U*_*d*_. Right: Overlaid snapshots of Yoda1 and Dooku binding poses, extracted from fully coupled (*λ* =1) trajectories of FEP/REMD.

#### FEP/REMD setup

The ABFEs were calculated with double-decoupling method^38^ using FEP/REMD implemented in NAMD2.14^37^. MD-based prediction of binding affinity at TM region requires accurate sampling of the ensemble distributions of ligand, water, protein, and lipids in bound and unbound states. To achieve this goal, we take advantage of the massive paralleled FEP/REMD^37^ to sample 128 intermediates between initial (unbound) and final (bound) end states. The soft-core potential was used to scale van der Walls interaction to prevent the occurrence of singularities at small *λ* values^55^. All simulations were conducted at 300 K and 1 bar NPT ensemble. PME is used for long-range electrostatics and short range cutoff is 12 Å with a smooth switching off between 10 to 12 Å. Long range correction cutoff is 25 Å (see below).

#### ABFE calculation protocol

To compute 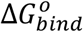, let’s set *U* as the total potential energy of the system (*λ*=1) and *U*_0_ is the total potential energy when the ligand is fully decoupled from its environment (*λ*=0). The dissociate constant is 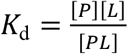, where [*p*], [*L*], [*pL*] are the equilibrium concentrations of ligand, protein, and bound complex, respectively. We define *L* as the coordinates of one Yoda1 molecule at low [*L*], *X* represents the coordinates of the solvent, lipids, protein. *r*_l_ is the center of mass position of Yoda1, and *r∗* is any location in the bulk region. In a homogenous and isotropic bulk, assume the number of ligands is larger than number of receptors, the dissociate constant can be expressed as the ratio of configurational integrals following the formulism of Deng and Roux^56^,

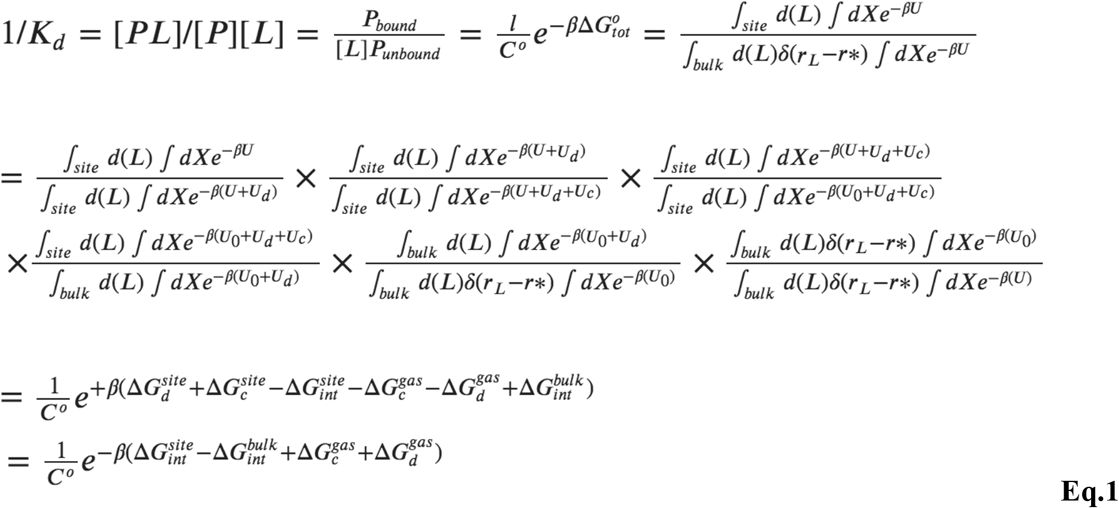

where the subscripts *site* and *bulk* indicate that the integrals include only configurations in which the ligand is in the binding site or bulk. The Dirac delta function δ(*r*_M_ *− r*^∗^) equals 0 if ligand is not in bulk (*r*_M_ ≠ *r*^*∗*^) and has a dimension of 1/volume. With the distance and conformational restraining potentials *U*_*c*_ and *U*_*d*_, the total reversible work to bring Yoda1 from bulk to binding site is decomposed into four sequential steps:

- The first and second integral ratios represent the free energy contribution of *U*_*c*_ and *U*_*d*_ at fully coupled state (*λ*=1), which are negligible based on our design of the flat-bottom harmonic restraints 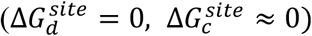
- The third integral ratio represents the free energy contribution of moving the ligand from gas phase to the binding site 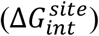, in presence of *U*, and *U*_*d*_ restraints. It is computed from FEP/REMD using 128 *λ* between 1*→*0.
- The fourth integral ratio is the restraint free energy from *U*_c,_ at fully decoupled state (i.e. in gas phase). It is computed from NAMD TI module with 20 equally spaced *λ* and 200 ps per *λ*. Each 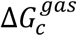 was computed three times and the average value was reported.
- The fifth integral ratio is the contribution of *U*_*d*_ when ligand is fully decoupled 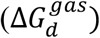, which is computed analytically as the ratio of volume sampled by flat-bottom distance restraint 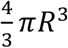 over standard molar concentration volume 1660 Å^3^.
- The last integral ratio represents the free energy contribution of decoupling the whole ligand in bulk 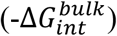, computed from FEP/REMD using 64 *λ* between 1*→*0.

With those restraints in place, a ligand is gradually decoupled from the binding site using 128 *λ* intermediates to obtain 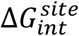, which is the free energy cost of moving the ligand from binding site to gas phase in presence of *R* and *RMSD* restraints. The *λ* =0 distribution (**Fig. 5** blue histograms) show that the ligand sampling is only restricted by the upper boundary of *RMSD* restraint potential *U*_*c*_, not the *R* restraint *U*_*d*_. The restraint free energy from *U*_*c*_ at fully decoupled (*λ* =0) state, 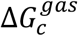, was computed using NAMD TI module and the average of three independent runs was reported. The contribution of *U*_*d*_, 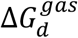 only factors in at the decoupled state after *U*_*c*_ is removed, thus can be computed analytically. The free energy contribution of decoupling the whole ligand in bulk 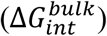 was obtained from FEP/REMD using 64 *λ* between 1*→*0, without restraint.

#### Long range corrections (LRC)

To account for the long range vdW interaction in nonisotropic binding environment beyond the CHARMM force field standard cutoff distance (12 Å with a smooth switching off between 10 to 12 Å), we recomputed the vdW potential energies using a larger vdW cutoff (25 Å with a smooth switching off between 23 to 25 Å) for every frame sampling in the fully coupled and fully decoupled states in bulk and in binding site. Specifically, we used the exponential averaging approach (also known as the Zwanzig relation)^57^, in which 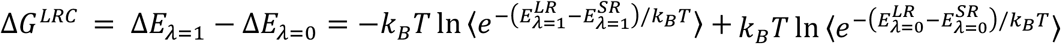. The average vdW energy ⟨*E*⟩ computed at different LJ cutoff distances from FEP/REMD simulations are plotted in **Fig. S6** and listed in **Table S2**. The ABFE results with and without LRCs are listed in **Table S1**. The LRCs contributed equally (−1.3 kcal/mol) to 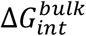 and 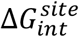 in closed state, thus do not change the binding affinity of Yoda1 and Dooku1 in the closed state. LRC’s contribution to 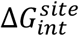 in open state is larger (−2.6 kcal/mol in Yoda1 and -2.1 kcal/mol in Dooku1), thus makes the binding affinity of Yoda1 and Dooku1 in the open state about 1 kcal/mol more favorable than the ABFE without LRC. Overall, the ABFE results with and without LCRs reach the same conclusion that Yoda1 binds to the open state stronger than the closed state, and Dooku1 binds to the closed state stronger than the open state.

#### Convergence of ABFE

Yoda1 or Dooku1 is gradually decoupled from the binding site (128 *λ* intermediates) and bulk solvent (64 *λ* intermediates). The time evolution of the FEP/REMD results suggest that a plateau is reached within 4 to 6 ns/replica for all six protein-ligand systems (**Fig S1**). We considered the FEP/REMD simulation approximates convergence when that last two 0.5 ns running average of free energy value fluctuates within 0.5 kcal/mol. Neighboring replica exchange was attempted at every 100 steps (0.2 ps) and yielded >70% acceptance ratios for all systems (**Fig. S2**). The numbers of uncorrelated sampling per replica are plotted in **Fig. S3**. Round trip time is about 2ns (**Fig. S4 and S5**). The convergence of *RMSD* restraint free energy at *λ* =0 is plotted in **Fig. S9**. The final results and uncertainties are summarized in **Table S1**.

#### Compare the end states of FEP/REMD with unbiased simulations

In order to validate whether our FEP simulations captured the water/lipid relationship inside the open and closed pocket, we defined the distance between a non-hydrogen atom and the center of mass of binding pocket residues (PDBID 6B3R: 1683-1733,2047-2097) and set 8Å as the cutoff distance to be inside the pocket. Results (**Fig. S9**) showed that there were ∼20 non-hydrogen lipid atoms inside the pocket and in the unbiased MD simulation in open state without Yoda1 and ∼25 in the FEP open state simulations at *λ* =0, which shows the similar magnitude for apo binding pocket. For the open pocket, there are about 1 or 2 water oxygen atoms inside in unbiased MD while shows the same case in FEP simulation at *λ* =0. For closed pocket, the similar trend was also observed that more water oxygen atoms inside the pocket about ∼5 for both unbiased and FEP simulations at *λ* =0 while there is almost no non-hydrogen lipid atoms inside the apo closed pocket. Overall, this atom number time series confirmed that our FEP simulations captured the water/lipid features inside the apo open and closed pockets, and thus ensure the accuracy of FEP samplings and calculations.

### Relative binding free energy simulations

The coordinates of the Piezo1 open and closed systems were taken from ABFE simulations and used as starting points for relative binding free energy (RBFE) calculations with thermodynamic integration (TI)^58^ method in AMBER. The AMBER force field (ff19SB^59^ for protein, GAFF2.1^60^ for ligand, and Lipid17 for lipids) was used for all the simulation systems. CPPTRAJ^61^, parmed^62^, and tleap were used to build the AMBER-TI systems. Each transformation pair system was generated with unified protocol in which both electrostatic and vdW interactions are scaled simultaneously by the softcore potentials^63-64^. The RBFE between two ligands (L0 to L1) is calculated as

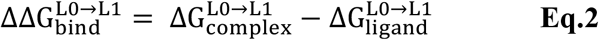

where 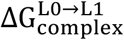 and 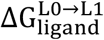 are the alchemical transformations of L0 to L1 in the complex and solution, respectively. The free energy difference between states L0 and L1 can be estimated as

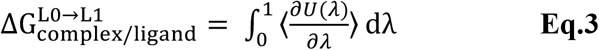

where U(*λ*) is the *λ*-coupled potential energy and *λ* is a coupling parameter varying from 0 to 1. The integration is calculated via the average of the *λ* derivative of the coupled potential energy at each intermediate *λ* state. The ΔG values are obtained by the sum of numerical integration over the number of *λ* windows quadrature points with associated weights of ∂U/ ∂*λ*. Long-range electrostatics was treated with the particle mesh Ewald (PME) method, and the van der Waals interactions were calculated with a cutoff distance of 10 Å ^65-66^. The second-order smoothstep softcore potential, SSC(2), was applied in the simulation^63^. The values of 0.2 and 50 Å^2^ were used for the parameters α and β of the softcore potential. Equilibration was performed for 5 ps employing the NPT ensemble after minimization in each *λ* window. AMBER-TI simulations were performed in the NPT ensemble at 310 K and 1 atm with the pmemd.cuda module of AMBER20 ^63^. Pressure was regulated by isotropic MC barostat with a pressure relaxation time of 2.0 ps. All alchemical transformations were done using the unified protocol with a 4-fs time step with hydrogen mass repartitioning (HMR)^67^.

#### Convergence of RBFE

In this study, 12 windows (0.0000, 0.0479, 0.1151, 0.2063, 0.3161, 0.4374, 0.5626, 0.6839, 0.7937, 0.8850, 0.9521, and 1.0000) were employed for each complex and solution system. 5 ns AMBER-TI simulations were performed for each *λ* window and the last 4 ns was used to calculate the final ΔΔG_bind_. For the transformation pair from Yoda1 to Dooku1 in the closed state, 10 ns AMBER-TI simulations were performed for each *λ* window to ensure the estimated ΔΔG_bind_ value is obtained from the converged trajectories. For this transformation pair, the last 5 ns of AMBER-TI simulations were used to estimate the ΔΔG_bind_. Four independent runs for each ΔΔG_bind_were performed for statistical analysis. Convergence plots for each transformation are provided in **Fig. S10-11**. For convergence analysis, the trajectories used for the final ΔΔG _bind_ were divided into 12 blocks to estimate the cumulative average ΔΔG_bind_ in the forward direction. The cumulative ΔΔG _bind_ values were obtained from the time-reversed data starting from the end of trajectories but using the same amount of simulation time^68^. The forward and time-reversed free energy estimations agree within error, indicating that our AMBER-TI simulations are converged well^69^.

### Site Identification by Ligand Competitive Saturation

Site Identification by Ligand Competitive Saturation (SILCS) was performed using the MolCal program (SilcsBio, LLC) and GROMACS 2016.4 simulation program^70^. The SILCS simulation uses an iterative Grand Canonical Monte Carlo (GCMC) and MD protocol, in which GCMC involves the sampling of water and various small molecules representing different functional groups, with the subsequent MD simulation for the sampling of protein conformational dynamics^71^. In this approach, benzene, propane, methanol, formamide, imidazole, acetaldehyde, methylammonium, and acetate in standard version and fluoroethane, trifluoroethane, chloroethane, fluorobenzene, chlorobenzene, bromobenzene, dimethylether, and methanol in halogen version were used as drug-like functional groups. During the simulations, these functional groups and water molecules compete with each other to bind with the protein. From ten copies of 100 ns trajectories, the three-dimensional (3D) probability distributions of the functional groups, called FragMaps, were calculated and then converted into a free energy representation, termed Grid Free Energy (GFE), based on a Boltzmann distribution for quantitative analysis. Force field for protein, water, ions and small molecules remains the same as in FEP calculations. The overlap coefficients are between 0.75 to 0.97 for all FragMaps obtained in current study.

The Monte Carlo based SILCS (SILCS-MC) protocol^42^ was applied to predict ligand binding poses using SILCS GFE as the scoring function. In this study, we utilized a full version of FragMaps with standard maps for generic apolar (benzene, propane C), generic H-bond donor (methanol O, formamide N, imidazole (NH)) and acceptor (methanol O, formamide O, imidazole N, acetaldehyde O), negatively charged (acetate) and positively charged (methylammonium) and halogen maps for fluoroethane fluorine (FETX), trifluoroethane carbon (TFEC), fluorobenzene fluorine (FLBX), chloroethane chlorine (CLEX), chlorobenzene chlorine (CLBX), bromobenzene bromine (BRBX), and dimethyl ether oxygen (DMEO) to supplement the 2018 generic halogen atom classification scheme (ACS) for scoring. With the improved treatment of halogens in CGenFF, halogen maps are used to reproduce the σ-holes and improve halogen bonding. For each docking, five independent SILCS-MC runs were performed with a two-step mode. Each SILCS-MC run involved a long-local MC protocol of 250 cycles of 10000 Metropolis MC steps and 40000 steps of simulated annealing (SA) from initial aligned structures. MC steps sample a wide range of binding poses with step-size of 1Å for translations, 180° for maximal rigid rotations per step and 180° for maximal dihedral angle rotations per step. This setup is to ensure sufficient sampling of ligand poses at the binding pocket. SA steps are designed to find a local minimum with the acceptance criteria defined by the Ligand Grid Free Energies (LGFE). SA involved maximum ranges of step-size of 0.2Å for translations, 9° for maximal rigid rotations per step and 9° for maximal dihedral angle rotations per step with gradual cooling from 300 to 0 K. The SILCS-MC runs were repeated for five times during which each cycle initiated with a different random seed and was repeated until it finished 250 cycles or reached convergence. The lowest LGFE and average LGFE scores from the minimum score in five independent jobs were obtained. The SILCS-MC exhaustive protocol was also tested from different initial ligand poses located around Pore Anchor (residue 2143 to 2175) or between Repeat A and Repeat B or between Repeat B and Repeat C, which shows the same setting as long-local MC protocol except ligand placement radius of 10Å.

## Acknowledgement

This work was supported by NIH grant GM130834 to Y.L.L. and J.J.L, and GM138472 to W.I.. Computational resources for ABFE calculations were provided by NSF XSEDE research allocation MCB160119 (Y.L.L.). Authors would like to thank the SilcsBio team, especially Prof. Alexander MacKerell and Dr. Sunhwan Jo for their support and suggestions.

## Notes

### Competing Interest Statement

The authors have declared no competing interest.

## References

1. Payandeh, J.; Volgraf, M., Ligand binding at the protein-lipid interface: strategic considerations for drug design. Nat Rev Drug Discov 2021, 20 (9), 710–722.

2. Zhao, Q.; Zhou, H.; Chi, S.; Wang, Y.; Wang, J.; Geng, J.; Wu, K.; Liu, W.; Zhang, T.; Dong, M.-Q.; Wang, J.; Li, X.; Xiao, B., Structure and mechanogating mechanism of the Piezo1 channel. Nature 2018.

3. Saotome, K.; Murthy, S. E.; Kefauver, J. M.; Whitwam, T.; Patapoutian, A.; Ward, A. B., Structure of the mechanically activated ion channel Piezo1. Nature 2018, 554 (7693), 481–486.

4. Guo, Y. R.; MacKinnon, R., Structure-based membrane dome mechanism for Piezo mechanosensitivity. Elife 2017, 6.

5. Ge, J.; Li, W.; Zhao, Q.; Li, N.; Chen, M.; Zhi, P.; Li, R.; Gao, N.; Xiao, B.; Yang, M., Architecture of the mammalian mechanosensitive Piezo1 channel. Nature 2015, 527 (7576), 64–9.

6. Wang, L.; Zhou, H.; Zhang, M.; Liu, W.; Deng, T.; Zhao, Q.; Li, Y.; Lei, J.; Li, X.; Xiao, B., Structure and mechanogating of the mammalian tactile channel PIEZO2. Nature 2019, 573 (7773), 225–229.

7. Syeda, R.; Xu, J.; Dubin, A. E.; Coste, B.; Mathur, J.; Huynh, T.; Matzen, J.; Lao, J.; Tully, D. C.; Engels, I. H.; Petrassi, H. M.; Schumacher, A. M.; Montal, M.; Bandell, M.; Patapoutian, A., Chemical activation of the mechanotransduction channel Piezo1. Elife 2015, 4.

8. Yang, Y.; Yao, K.; Repasky, M. P.; Leswing, K.; Abel, R.; Shoichet, B. K.; Jerome, S. V., Efficient Exploration of Chemical Space with Docking and Deep Learning. J Chem Theory Comput 2021, 17 (11), 7106–7119.

9. Passini, F. S.; Jaeger, P. K.; Saab, A. S.; Hanlon, S.; Chittim, N. A.; Arlt, M. J.; Ferrari, K. D.; Haenni, D.; Caprara, S.; Bollhalder, M.; Niederost, B.; Horvath, A. N.; Gotschi, T.; Ma, S.; Passini-Tall, B.; Fucentese, S. F.; Blache, U.; Silvan, U.; Weber, B.; Silbernagel, K. G.; Snedeker, J. G., Shear-stress sensing by PIEZO1 regulates tendon stiffness in rodents and influences jumping performance in humans. Nat Biomed Eng 2021.

10. Deivasikamani, V.; Dhayalan, S.; Abudushalamu, Y.; Mughal, R.; Visnagri, A.; Cuthbertson, K.; Scragg, J. L.; Munsey, T. S.; Viswambharan, H.; Muraki, K.; Foster, R.; Sivaprasadarao, A.; Kearney, M. T.; Beech, D. J.; Sukumar, P., Piezo1 channel activation mimics high glucose as a stimulator of insulin release. Sci Rep 2019, 9 (1), 16876.

11. Caolo, V.; Debant, M.; Endesh, N.; Futers, T. S.; Lichtenstein, L.; Bartoli, F.; Parsonage, G.; Jones, E. A.; Beech, D. J., Shear stress activates ADAM10 sheddase to regulate Notch1 via the Piezo1 force sensor in endothelial cells. Elife 2020, 9.

12. Morley, L. C.; Shi, J.; Gaunt, H. J.; Hyman, A. J.; Webster, P. J.; Williams, C.; Forbes, K.; Walker, J. J.; Simpson, N. A. B.; Beech, D. J., Piezo1 channels are mechanosensors in human fetoplacental endothelial cells. Mol Hum Reprod 2018, 24 (10), 510–520.

13. Cahalan, S. M.; Lukacs, V.; Ranade, S. S.; Chien, S.; Bandell, M.; Patapoutian, A., Piezo1 links mechanical forces to red blood cell volume. Elife 2015, 4.

14. Jesse R. Holt; Wei-Zheng Zeng; Elizabeth L. Evans; Seung-Hyun Woo; Shang Ma; Hamid Abuwarda; Meaghan Loud; Ardem Patapoutian; Pathak, M. M., Spatiotemporal dynamics of PIEZO1 localization controls keratinocyte migration during wound healing. BioRxiv 2020.

15. Bosutti, A.; Giniatullin, A.; Odnoshivkina, Y.; Giudice, L.; Malm, T.; Sciancalepore, M.; Giniatullin, R.; D’Andrea, P.; Lorenzon, P.; Bernareggi, A., “Time window” effect of Yoda1-evoked Piezo1 channel activity during mouse skeletal muscle differentiation. Acta Physiol (Oxf) 2021, e13702.

16. Jetta, D.; Gottlieb, P. A.; Verma, D.; Sachs, F.; Hua, S. Z., Shear stress-induced nuclear shrinkage through activation of Piezo1 channels in epithelial cells. J Cell Sci 2019, 132 (11).

17. Choi, D.; Park, E.; Jung, E.; Cha, B.; Lee, S.; Yu, J.; Kim, P. M.; Lee, S.; Hong, Y. J.; Koh, C. J.; Cho, C. W.; Wu, Y.; Li Jeon, N.; Wong, A. K.; Shin, L.; Kumar, S. R.; Bermejo-Moreno, I.; Srinivasan, R. S.; Cho, I. T.; Hong, Y. K., Piezo1 incorporates mechanical force signals into the genetic program that governs lymphatic valve development and maintenance. JCI Insight 2019, 4 (5).

18. Romac, J. M.; Shahid, R. A.; Swain, S. M.; Vigna, S. R.; Liddle, R. A., Piezo1 is a mechanically activated ion channel and mediates pressure induced pancreatitis. Nat Commun 2018, 9 (1), 1715.

19. Lacroix, J. J.; Botello-Smith, W. M.; Luo, Y., Probing the gating mechanism of the mechanosensitive channel Piezo1 with the small molecule Yoda1. Nature Communications 2018, 9 (1), 2029.

20. Nonomura, K.; Lukacs, V.; Sweet, D. T.; Goddard, L. M.; Kanie, A.; Whitwam, T.; Ranade, S. S.; Fujimori, T.; Kahn, M. L.; Patapoutian, A., Mechanically activated ion channel PIEZO1 is required for lymphatic valve formation. Proc Natl Acad Sci U S A 2018.

21. Wang, S.; Chennupati, R.; Kaur, H.; Iring, A.; Wettschureck, N.; Offermanns, S., Endothelial cation channel PIEZO1 controls blood pressure by mediating flow-induced ATP release. The Journal of clinical investigation 2016, 126 (12), 4527–4536.

22. Beech, D. J., Endothelial Piezo1 channels as sensors of exercise. J Physiol 2018, 596 (6), 979–984.

23. Nguetse, C. N.; Purington, N.; Ebel, E. R.; Shakya, B.; Tetard, M.; Kremsner, P. G.; Velavan, T. P.; Egan, E. S., A common polymorphism in the mechanosensitive ion channel PIEZO1 is associated with protection from severe malaria in humans. Proc Natl Acad Sci U S A 2020, 117 (16), 9074–9081.

24. Ma, S.; Cahalan, S.; LaMonte, G.; Grubaugh, N. D.; Zeng, W.; Murthy, S. E.; Paytas, E.; Gamini, R.; Lukacs, V.; Whitwam, T.; Loud, M.; Lohia, R.; Berry, L.; Khan, S. M.; Janse, C. J.; Bandell, M.; Schmedt, C.; Wengelnik, K.; Su, A. I.; Honore, E.; Winzeler, E. A.; Andersen, K. G.; Patapoutian, A., Common PIEZO1 Allele in African Populations Causes RBC Dehydration and Attenuates Plasmodium Infection. Cell 2018, 173 (2), 443–455 e12.

25. Sun, W. J.; Chi, S. P.; Li, Y. H.; Ling, S. K.; Tan, Y. J.; Xu, Y. J.; Jiang, F.; Li, J. W.; Liu, C. Z.; Zhong, G. H.; Cao, D. C.; Jin, X. Y.; Zhao, D. S.; Gao, X. C.; Liu, Z. Z.; Xiao, B. L.; Li, Y. X., The mechanosensitive Piezo1 channel is required for bone formation. Elife 2019, 8.

26. Evans, E. L.; Cuthbertson, K.; Endesh, N.; Rode, B.; Blythe, N. M.; Hyman, A. J.; Hall, S. J.; Gaunt, H. J.; Ludlow, M. J.; Foster, R.; Beech, D. J., Yoda1 analogue (Dooku1) which antagonizes Yoda1-evoked activation of Piezo1 and aortic relaxation. Br J Pharmacol 2018, 175 (10), 1744–1759.

27. Li, H.; Xiong, J.; Yan, W.; O’brien, C.; Schuller De Almeida, M. Piezo1 agonists for the promotion of bone formation. World Patent WO2021067943 A1. 2021.

28. Wijerathne, T. D.; Ozkan, A. D.; Lacroix, J. J., Yoda1’s energetic footprint on Piezo1 channels and its modulation by voltage and temperature. Proceedings of the National Academy of Sciences of the United States of America 2022, in Press.

29. Wang, Y.; Chi, S.; Guo, H.; Li, G.; Wang, L.; Zhao, Q.; Rao, Y.; Zu, L.; He, W.; Xiao, B., A lever-like transduction pathway for long-distance chemical-and mechano-gating of the mechanosensitive Piezo1 channel. Nat Commun 2018, 9 (1), 1300.

30. Botello-Smith, W. M.; Jiang, W.; Zhang, H.; Ozkan, A. D.; Lin, Y. C.; Pham, C. N.; Lacroix, J. J.; Luo, Y., A mechanism for the activation of the mechanosensitive Piezo1 channel by the small molecule Yoda1. Nature Communications 2019, 10 (1), 4503.

31. Lin, Y. C.; Guo, Y. R.; Miyagi, A.; Levring, J.; MacKinnon, R.; Scheuring, S., Force-induced conformational changes in PIEZO1. Nature 2019, 573 (7773), 230–234.

32. De Vecchis, D.; Beech, D. J.; Kalli, A. C., Molecular dynamics simulations of Piezo1 channel opening by increases in membrane tension. Biophys J 2021, 120 (8), 1510–1521.

33. Jiang, W.; Del Rosario, J. S.; Botello-Smith, W.; Zhao, S.; Lin, Y. C.; Zhang, H.; Lacroix, J.; Rohacs, T.; Luo, Y. L., Crowding-induced opening of the mechanosensitive Piezo1 channel in silico. Communications Biology 2021, 4 (1), 84.

34. Cox, C. D.; Bae, C.; Ziegler, L.; Hartley, S.; Nikolova-Krstevski, V.; Rohde, P. R.; Ng, C. A.; Sachs, F.; Gottlieb, P. A.; Martinac, B., Removal of the mechanoprotective influence of the cytoskeleton reveals PIEZO1 is gated by bilayer tension. Nat Commun 2016, 7, 10366.

35. Lewis, A. H.; Grandl, J., Mechanical sensitivity of Piezo1 ion channels can be tuned by cellular membrane tension. Elife 2015, 4.

36. Yang, X.; Lin, C.; Chen, X.; Li, S.; Li, X.; Xiao, B., Structure deformation and curvature sensing of PIEZO1 in lipid membranes. Nature 2022, 604 (7905), 377–383.

37. Jiang, W.; Hodoscek, M.; Roux, B., Computation of Absolute Hydration and Binding Free Energy with Free Energy Perturbation Distributed Replica-Exchange Molecular Dynamics. Journal of Chemical Theory and Computation 2009, 5 (10), 2583–2588.

38. Gilson, M. K.; Given, J. A.; Bush, B. L.; McCammon, J. A., The statistical-thermodynamic basis for computation of binding affinities: A critical review. Biophysical Journal 1997, 72 (3), 1047–1069.

39. Lacroix, J. J.; Botello-Smith, W. M.; Luo, Y., Probing the gating mechanism of the mechanosensitive channel Piezo1 with the small molecule Yoda1. Nat Commun 2018, 9 (1), 2029.

40. Tang, H.; Zeng, R.; He, E.; Zhang, I.; Ding, C.; Zhang, A., Piezo-Type Mechanosensitive Ion Channel Component 1 (Piezo1): A Promising Therapeutic Target and Its Modulators. J Med Chem 2022, 65 (9), 6441–6453.

41. Mousaei, M.; Kudaibergenova, M.; MacKerell, A. D., Jr.; Noskov, S., Assessing hERG1 Blockade from Bayesian Machine-Learning-Optimized Site Identification by Ligand Competitive Saturation Simulations. J Chem Inf Model 2020, 60 (12), 6489–6501.

42. Faller, C. E.; Raman, E. P.; MacKerell, A. D., Jr.; Guvench, O., Site Identification by Ligand Competitive Saturation (SILCS) simulations for fragment-based drug design. Methods Mol Biol 2015, 1289, 75–87.

43. Inc., C. C. G. Molecular Operating Environment (MOE), 2013.08; Montreal, QC, Canada, 2016.

44. Lee, J.; Cheng, X.; Swails, J. M.; Yeom, M. S.; Eastman, P. K.; Lemkul, J. A.; Wei, S.; Buckner, J.; Jeong, J. C.; Qi, Y.; Jo, S.; Pande, V. S.; Case, D. A.; Brooks, C. L., 3rd; MacKerell, A. D., Jr.; Klauda, J. B.; Im, W., CHARMM-GUI Input Generator for NAMD, GROMACS, AMBER, OpenMM, and CHARMM/OpenMM Simulations Using the CHARMM36 Additive Force Field. J Chem Theory Comput 2016, 12 (1), 405–13.

45. Jo, S.; Kim, T.; Iyer, V. G.; Im, W., CHARMM-GUI: a web-based graphical user interface for CHARMM. J Comput Chem 2008, 29 (11), 1859–65.

46. Mackerell Jr, A. D.; Feig, M.; Brooks III, C. L., Extending the treatment of backbone energetics in protein force fields: limitations of gas‐phase quantum mechanics in reproducing protein conformational distributions in molecular dynamics simulations. Journal of computational chemistry 2004, 25 (11), 1400–1415.

47. MacKerell Jr, A. D.; Bashford, D.; Bellott, M.; Dunbrack Jr, R. L.; Evanseck, J. D.; Field, M. J.; Fischer, S.; Gao, J.; Guo, H.; Ha, S., All-atom empirical potential for molecular modeling and dynamics studies of proteins. The journal of physical chemistry B 1998, 102 (18), 3586–3616.

48. Klauda, J. B.; Venable, R. M.; Freites, J. A.; O’Connor, J. W.; Tobias, D. J.; Mondragon-Ramirez, C.; Vorobyov, I.; MacKerell Jr, A. D.; Pastor, R. W., Update of the CHARMM all-atom additive force field for lipids: validation on six lipid types. The journal of physical chemistry B 2010, 114 (23), 7830–7843.

49. Jorgensen, W. L.; Chandrasekhar, J.; Madura, J. D.; Impey, R. W.; Klein, M. L., Comparison of simple potential functions for simulating liquid water. The Journal of chemical physics 1983, 79 (2), 926–935.

50. Vanommeslaeghe, K.; Hatcher, E.; Acharya, C.; Kundu, S.; Zhong, S.; Shim, J.; Darian, E.; Guvench, O.; Lopes, P.; Vorobyov, I., CHARMM general force field: A force field for drug‐like molecules compatible with the CHARMM all‐atom additive biological force fields. Journal of computational chemistry 2010, 31 (4), 671–690.

51. Yu, W.; He, X.; Vanommeslaeghe, K.; MacKerell Jr, A. D., Extension of the CHARMM general force field to sulfonyl‐containing compounds and its utility in biomolecular simulations. Journal of computational chemistry 2012, 33 (31), 2451–2468.

52. Mayne, C. G.; Saam, J.; Schulten, K.; Tajkhorshid, E.; Gumbart, J. C., Rapid parameterization of small molecules using the force field toolkit. J. Comput. Chem. 2013, 34 (32), 2757–2770.

53. Humphrey, W.; Dalke, A.; Schulten, K., VMD: visual molecular dynamics. J Mol Graph 1996, 14 (1), 33-8, 27-8.

54. Salari, R.; Joseph, T.; Lohia, R.; Hénin, J.; Brannigan, G., A streamlined, general approach for computing ligand binding free energies and its application to GPCR-bound cholesterol. Journal of chemical theory and computation 2018, 14 (12), 6560–6573.

55. Zacharias, M.; Straatsma, T. P.; Mccammon, J. A., Separation-Shifted Scaling, a New Scaling Method for Lennard-Jones Interactions in Thermodynamic Integration. J Chem Phys 1994, 100 (12), 9025–9031.

56. Deng, Y. Q.; Roux, B., Calculation of standard binding free energies: Aromatic molecules in the T4 lysozyme L99A mutant. Journal of Chemical Theory and Computation 2006, 2 (5), 1255–1273.

57. Shirts, M. R.; Mobley, D. L.; Chodera, J. D.; Pande, V. S., Accurate and efficient corrections for missing dispersion interactions in molecular simulations. J Phys Chem B 2007, 111 (45), 13052–63.

58. Kumar, J.; Dey, T. K.; Sinha, S. K., Semiclassical statistical mechanics of hard-body fluid mixtures. J Chem Phys 2005, 122 (22), 224504.

59. Tian, C.; Kasavajhala, K.; Belfon, K. A. A.; Raguette, L.; Huang, H.; Migues, A. N.; Bickel, J.; Wang, Y.; Pincay, J.; Wu, Q.; Simmerling, C., ff19SB: Amino-Acid-Specific Protein Backbone Parameters Trained against Quantum Mechanics Energy Surfaces in Solution. J Chem Theory Comput 2020, 16 (1), 528–552.

60. He, X.; Man, V. H.; Yang, W.; Lee, T. S.; Wang, J., A fast and high-quality charge model for the next generation general AMBER force field. J Chem Phys 2020, 153 (11), 114502.

61. Roe, D. R.; Cheatham, T. E., 3rd, PTRAJ and CPPTRAJ: Software for Processing and Analysis of Molecular Dynamics Trajectory Data. J Chem Theory Comput 2013, 9 (7), 3084–95.

62. Shirts, M. R.; Klein, C.; Swails, J. M.; Yin, J.; Gilson, M. K.; Mobley, D. L.; Case, D. A.; Zhong, E. D., Lessons learned from comparing molecular dynamics engines on the SAMPL5 dataset. J Comput Aided Mol Des 2017, 31 (1), 147–161.

63. Lee, T. S.; Allen, B. K.; Giese, T. J.; Guo, Z.; Li, P.; Lin, C.; McGee, T. D., Jr.; Pearlman, D. A.; Radak, B. K.; Tao, Y.; Tsai, H. C.; Xu, H.; Sherman, W.; York, D. M., Alchemical Binding Free Energy Calculations in AMBER20: Advances and Best Practices for Drug Discovery. J Chem Inf Model 2020, 60 (11), 5595–5623.

64. Zhang, H.; Kim, S.; Giese, T. J.; Lee, T. S.; Lee, J.; York, D. M.; Im, W., CHARMM-GUI Free Energy Calculator for Practical Ligand Binding Free Energy Simulations with AMBER. J Chem Inf Model 2021, 61 (9), 4145–4151.

65. Darden, T.; York, D.; Pedersen, L., Particle Mesh Ewald - an N.Log(N) Method for Ewald Sums in Large Systems. J Chem Phys 1993, 98 (12), 10089–10092.

66. Essmann, U.; Perera, L.; Berkowitz, M. L.; Darden, T.; Lee, H.; Pedersen, L. G., A Smooth Particle Mesh Ewald Method. J Chem Phys 1995, 103 (19), 8577–8593.

67. Hopkins, C. W.; Le Grand, S.; Walker, R. C.; Roitberg, A. E., Long-Time-Step Molecular Dynamics through Hydrogen Mass Repartitioning. Journal of Chemical Theory and Computation 2015, 11 (4), 1864–1874.

68. Yang, W.; Bitetti-Putzer, R.; Karplus, M., Free energy simulations: Use of reverse cumulative averaging to determine the equilibrated region and the time required for convergence. J Chem Phys 2004, 120 (6), 2618–2628.

69. Klimovich, P. V.; Shirts, M. R.; Mobley, D. L., Guidelines for the analysis of free energy calculations. J Comput Aid Mol Des 2015, 29 (5), 397–411.

70. Hess, B.; Kutzner, C.; van der Spoel, D.; Lindahl, E., GROMACS 4: Algorithms for highly efficient, load-balanced, and scalable molecular simulation. Journal of Chemical Theory and Computation 2008, 4 (3), 435–447.

71. Lakkaraju, S. K.; Raman, E. P.; Yu, W.; MacKerell, A. D., Jr., Sampling of Organic Solutes in Aqueous and Heterogeneous Environments Using Oscillating Excess Chemical Potentials in Grand Canonical-like Monte Carlo-Molecular Dynamics Simulations. J Chem Theory Comput 2014, 10 (6), 2281–2290.

