## Supplemental material for "Binding Free Energies of Piezo1 Channel Agonists at Protein-Membrane Interface"

### Supplementary MD simulation protocols

For each system, a total of 7500 cycles of minimization were run, with 5000 steepest decent cycles followed by 2500 conjugate gradient cycles. For this stage and all following, a non-bonded cut-off of 12.0 Å plus a 10.0 Å force switching range was employed. Both positional and internal conformational restraints were applied. Positional restraints were applied to all atoms of all protein residues with a 10.0 kcal/mol/Å harmonic spring constant while phosphate atoms of membrane residues were restrained with a force constant of 2.5 kcal mol$^{-1}$ Å$^{-1}$. During minimization dihedral restraints were also employed to ensure retention of backbone phi-psi angles with 250 kcal mol$^{-1}$ radian$^{-2}$ force constant. Following minimization, a 25 ps NVT solvent heating simulation was run to obtain a target temperature of 310 K. For this stage through the fifth equilibration stage, a time step of 1 fs was employed. The same positional and internal restraint setup was applied as the previous minimization stage. For all simulation stages, temperature control was accomplished using a Langevin thermostat with a gamma parameter (friction coefficient) of 1.0 ps$^{-1}$. The SHAKE algorithm was used to constrain bonds involving hydrogens. The second equilibration stage followed an identical setup with the sole exception that the force constant of the protein positional restraints was reduced to 5.0 kcal mol$^{-1}$ Å$^{-1}$. The force constant of the dihedral phi-psi restraint was also reduced to 100 kcal mol$^{-1}$ radian$^{-2}$. The first and second stage equilibrations were each run for a duration of 25 ps each. Starting with the third equilibration stage, the duration of simulations was raised to 100 ps and an NPT ensemble was employed, with pressure regulation accomplished by utilization of a semi-isotropic Monte-Carlo barostat with a target pressure of 1.0 bar and constant zero membrane surface tension. For the third stage, force constants for protein and membrane positional restraints were reduced to 2.5 and 1.0 kcal mol$^{-1}$ Å$^{-1}$, respectively. The dihedral phi-psi protein backbone restraints were also dropped to 50.0 kcal mol$^{-1}$ radian$^{-2}$. In the fourth stage, positional restraint force constants for protein and membrane were reduced to 1.0 and 0.5 kcal mol$^{-1}$ Å$^{-1}$, respectively. In the fifth stage, positional restraint force constants for protein and membrane were reduced to 0.5 and 0.1 kcal mol$^{-1}$ Å$^{-1}$and phi-psi backbone dihedral restraint force constants were reduced to 25.0 kcal mol$^{-1}$ radian$^{-2}$. For the sixth and last equilibration simulation, membrane positional restraints and phi-psi backbone dihedral restraints were removed entirely while protein positional restraints were reduced to 0.1 kcal mol$^{-1}$ Å$^{-1}$. Simulation timestep was also raised to 2.0 fs as it would remain for all future simulations. In the end, an equilibration production run with only weak RMSD restraints for the first and last helices of the single arm was conducted for 5 replicas with different velocities and each for 100ns (all other parameters were identical to the sixth stage of equilibration).

**Table S1. Absolute binding free energy (ABFE) results. Data with long-range corrections are shown in parentheses.**

| Free energy kcal/mol | **Yoda-closed** | **Yoda-open** | **Yoda-open-switch** | **Dooku-closed** | **Dooku-open** | **Dooku-open-switch** |
|---|---|---|---|---|---|---|
| $\Delta G_{int}^{site}$ | -21.6 ±0.02 (-23.0 ±0.14) | -25.0 ±0.02 (-27.6 ±0.14) | -23.6 ±0.02 | -22.4 ±0.03 (-23.7 ±0.14) | -22.3 ±0.03 (-24.4 ±0.11) | -23.3 ±0.03 |
| $\Delta G_{int}^{bulk}$ | -11.4 ±0.02 (-12.8 ±0.10) | | | -10.6 ±0.05 (-11.9 ±0.11) | | |
| $\Delta G_{c\ \ \lambda=0}^{gas}$ | 4.7 ±0.2 | 4.7 ±0.1 | 4.3 ±0.6 | 3.4 ±0.6 | 5.4 ±0.2 | 5.9 ±0.2 |
| $\Delta G_{c\ \ \lambda=1}^{gas}$ | - | - | - | - | - | 0.5 ±0.1 |
| $\Delta G_{d}^{gas}$ | -1.4 | -0.5 | -1.0 | -1.4 | -1.0 | -0.5 |
| $\Delta G_{bind}$ | -6.9 ±0.2 (-6.8 ±0.2) | -9.4 ±0.1 (-10.5 ±0.2) | -8.9 ±0.8 | -9.8 ±0.6 (-9.7 ±0.6) | -7.3 ±0.2 (-8.1 ±0.2) | -6.8 ±0.2 |
| $\Delta\Delta \boldsymbol{G}_{bind}^{o-c}$ | -2.5 ±0.2 (-3.7 ±0.3) | | | 2.5 ±0.6 (1.6 ±0.6) | | |

* $\Delta G_{bind} = \Delta G_{int}^{site} - \Delta G_{int}^{bulk} + \Delta G_{c}^{gas} + \Delta G_{d}^{gas}$. The standard deviations for $\Delta G_{int}^{site}$ and $\Delta G_{int}^{bulk}$ were calculated from MBAR. Restraint $\Delta G_{c}^{gas}$was calculated using three independent simulations. $\Delta G_{d}^{gas}$ is using the restraint constants regarding each system (see Methods).

**Table S2. The long-range correction for absolute binding energy calculation.**

| Energy type* (kcal/mol) | System type | | | | | |
|---|---|---|---|---|---|---|
| | Yoda-closed | Yoda-open | Dooku-closed | Dooku-open | Yoda-solv | Dooku-solv |
| $\langle E_{\lambda=1}^{LR} \rangle$ | 23788.5±18.1 | 35893.0±24.1 | 26373.9±18.3 | 23871.6±24.8 | 1635.7±32.5 | 1557.4±42.4 |
| $\langle E_{\lambda=1}^{SR} \rangle$ | 26347.6±18.1 | 39627.1±24.1 | 29046.3±18.3 | 26440.8±24.8 | 1682.1±32.5 | 1599.6±42.4 |
| $\boldsymbol{\Delta E_{\lambda=1}}$ | -2564.6±0.20 | -3741.1±0.21 | -2677.0±0.15 | -2574.6±0.24 | -46.4±0.03 | -42.3±0.03 |
| $\langle E_{\lambda=0}^{LR} \rangle$ | 23835.7±18.1 | 35903.2±39.6 | 26380.0±18.4 | 23913.8±24.2 | 1516.8±4.0 | 1368.8±3.7 |
| $\langle E_{\lambda=0}^{SR} \rangle$ | 26393.6±18.1 | 39636.6±39.6 | 29051.1±18.4 | 26481.6±24.2 | 1561.7±4.0 | 1409.7±3.7 |
| $\boldsymbol{\Delta E_{\lambda=0}}$ | -2562.7±0.17 | -3738.2±0.34 | -2676.2±0.17 | -2572.3±0.22 | -45.0±0.03 | -41.0±0.02 |
| $\boldsymbol{\Delta G^{LRC}}$ | -1.99±0.24 | -2.81±0.30 | -0.80±0.23 | -2.29±0.33 | -1.43±0.04 | -1.33±0.03 |

*The energy at cutoff of 12 Å represents $E_{\lambda=1}^{SR}$ and cutoff of 25 Å represents $E_{\lambda=1}^{LR}$. The long-range correction (LRC) of the computed free energy is then calculated using the exponential averaging approach (also known as the Zwanzig relation): $\Delta G^{LRC} = \Delta E_{\lambda=1} - \Delta E_{\lambda=0} = -k_B T \ln \langle e^{-(E_{\lambda=1}^{LR} - E_{\lambda=1}^{SR})/k_B T} \rangle + k_B T \ln \langle e^{-(E_{\lambda=0}^{LR} - E_{\lambda=0}^{SR})/k_B T} \rangle$.

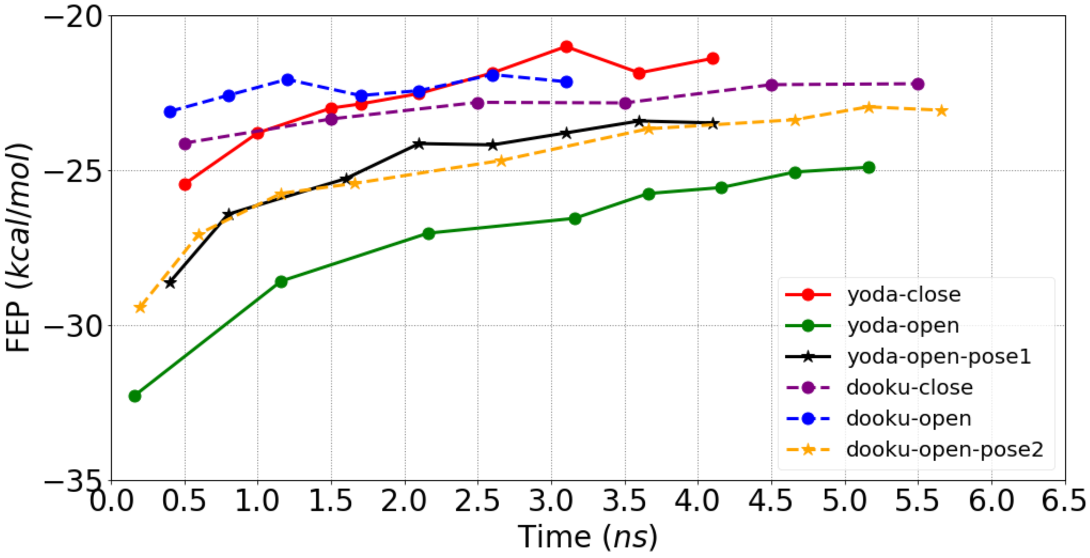


**Figure S1**. The convergence plot for all binding site ($\Delta G_{int}^{site}$) FEP/REMD systems labelled in different colors and markers. Yoda-open-pose1 and dooku-open-pose2 labels represent the open state system with Yoda1 and Dooku1 binding poses switched.

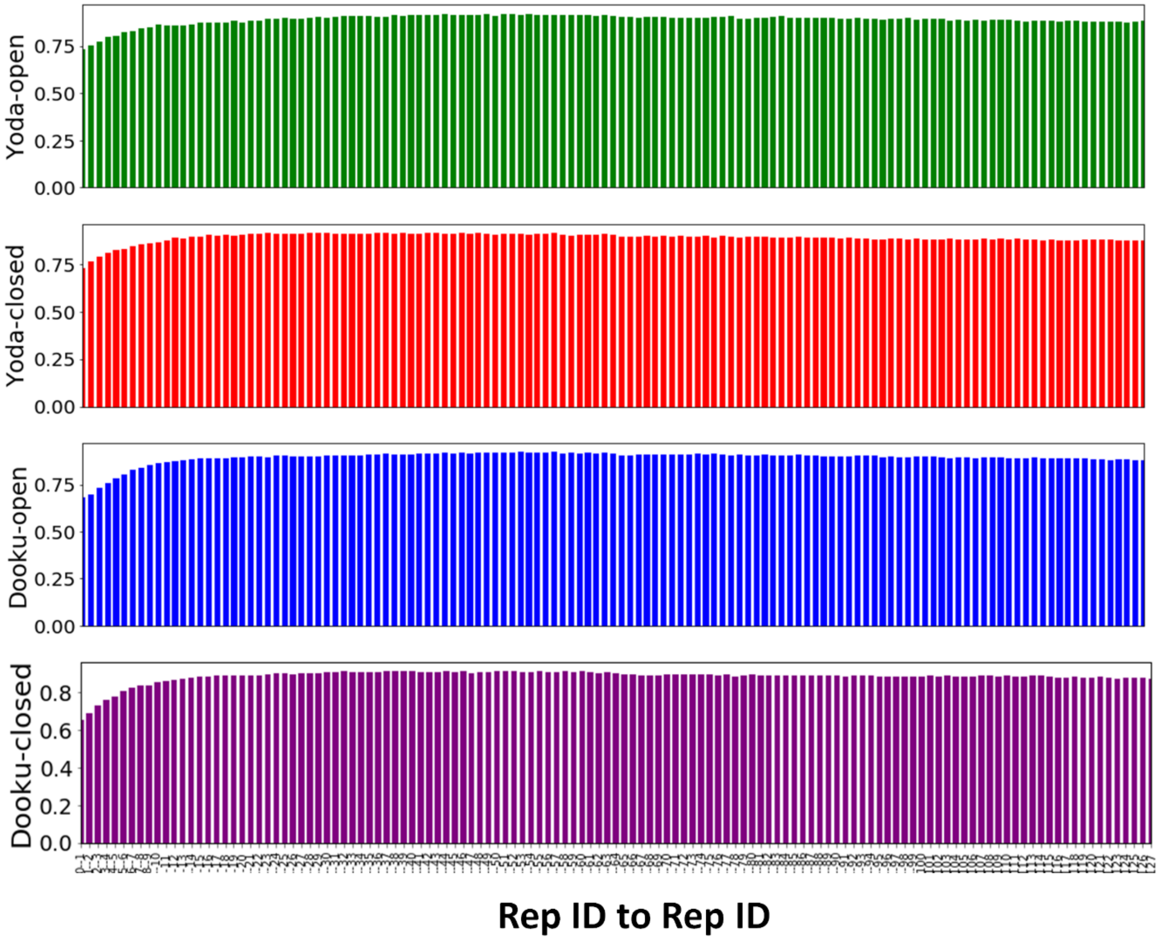


**Figure S2**. Acceptance ratio for each binding site ($\Delta G_{int}^{site}$) FEP/REMD system.

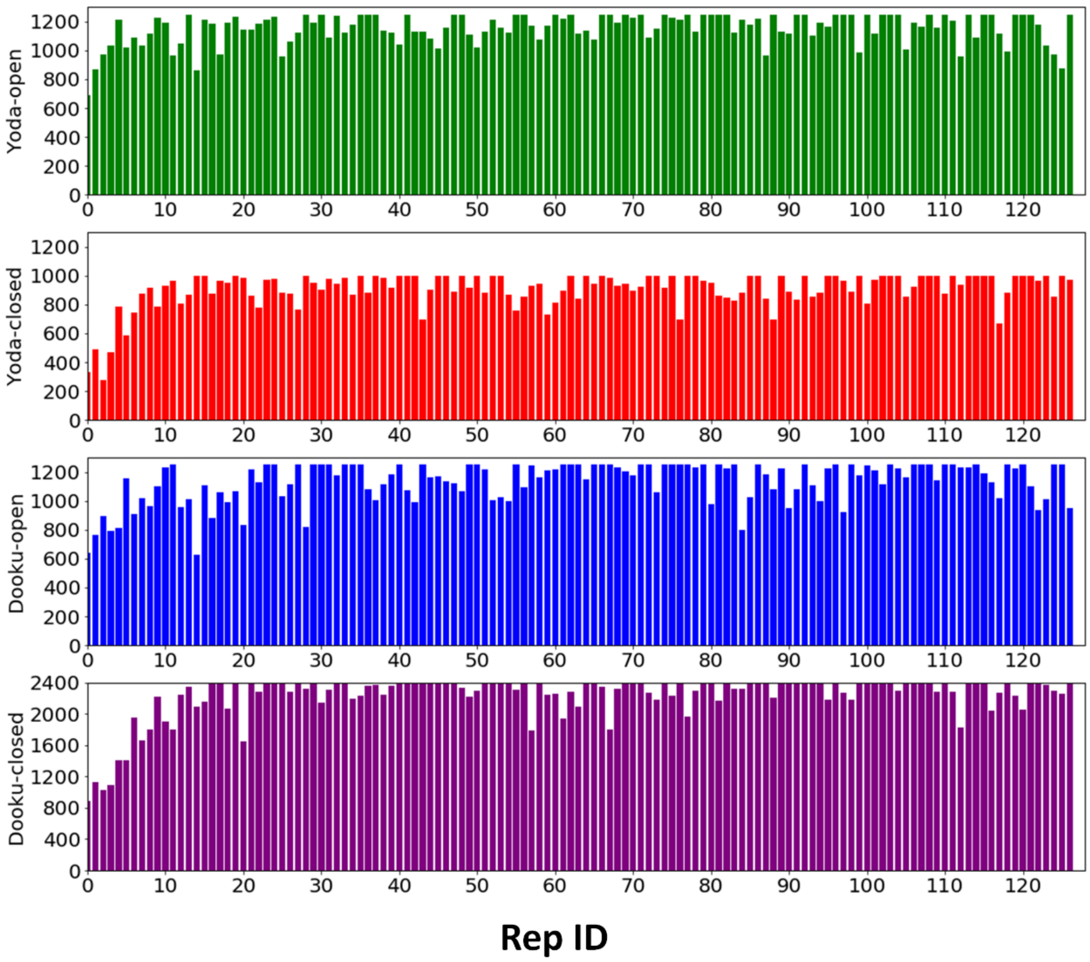


**Figure S3**. The number of uncorrelated samples in each replica for each binding site ($\Delta G_{int}^{site}$) FEP/REMD system.

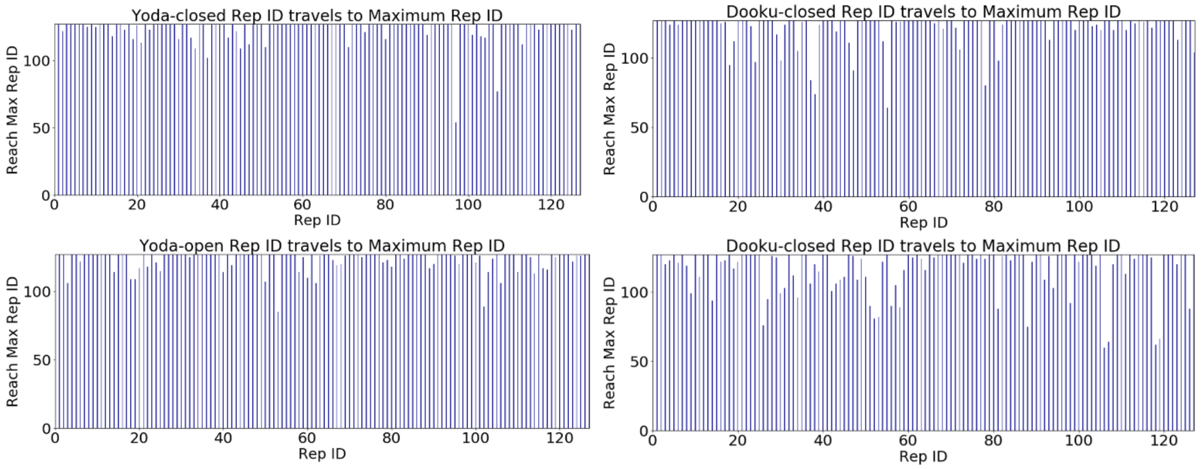


**Figure S4**. The number of replicas traveled during FEP/REMD. Rep ID in $x$-axis is the starting replica.

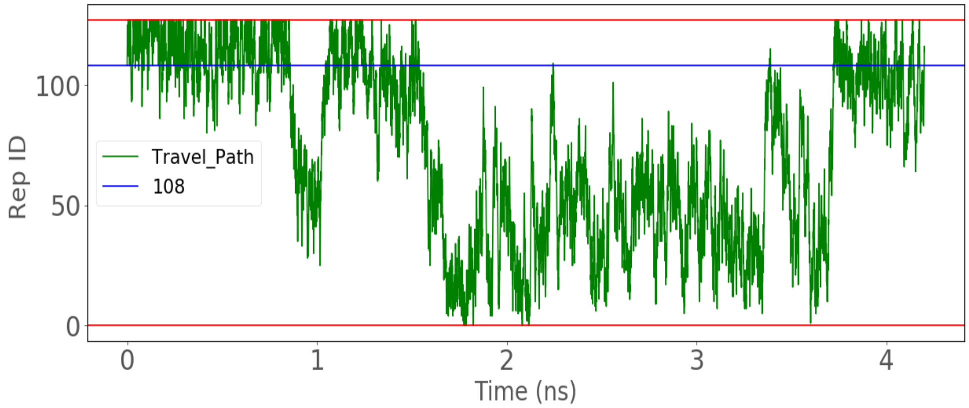


**Figure S5**. An example of the FEP/REMD replica travel history in Yoda1-closed state. Blue is the starting replica ID.

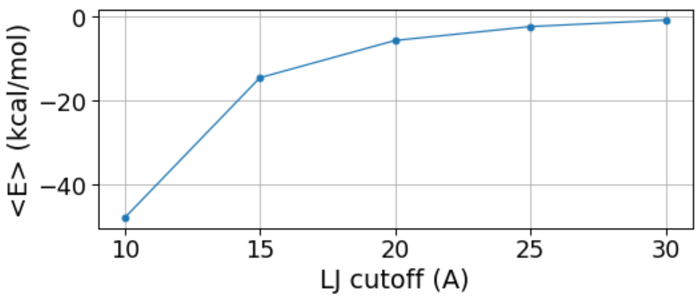


**Figure S6**. The difference in average vdW energy $\langle E \rangle$ computed between each LJ cutoff distance and the largest cutoff distance tested (30 Å) from FEP/REMD simulation of Yoda1 in solution ($\lambda = 1$), averaged over 500 snapshots.

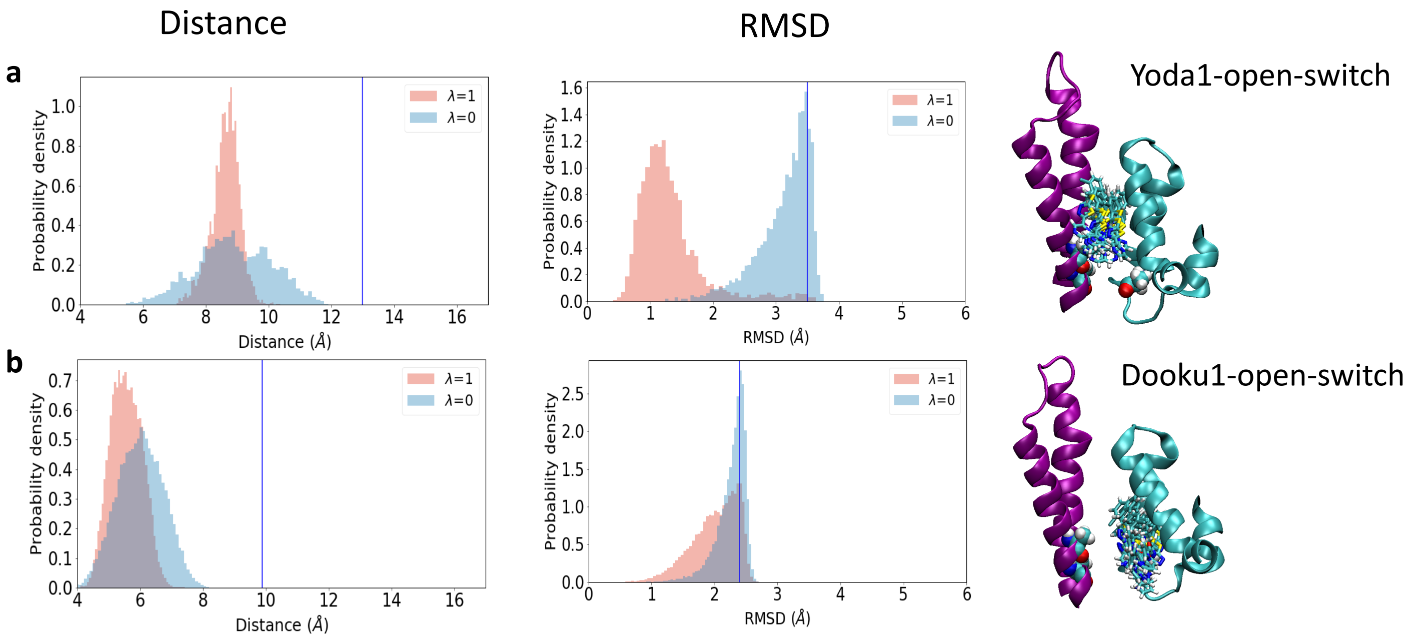


**Figure S7**. **FEP/REMD of switched binding poses in Piezo1 open state.** a) Distance R and RMSD distributions for Yoda1. b) Distance R and RMSD distributions for Dooku1. Blue vertical lines show the upper boundary of the distance R and RMSD restraints. Because the upper boundary of RMSD in Dooku1-open-switch biased the sampling of $\lambda$ =1, it's free energy contribution ($\Delta G_c^{gas}{}_{\lambda=1}$) was computed using NAMD TI module and listed in Table S1.

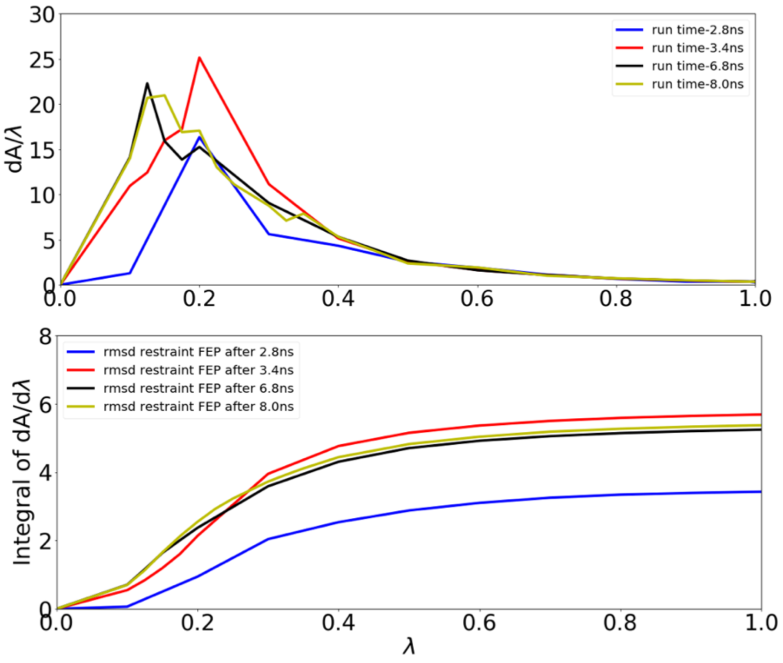


**Figure S8**. The convergence of RMSD restraint TI at $\lambda$ =0.

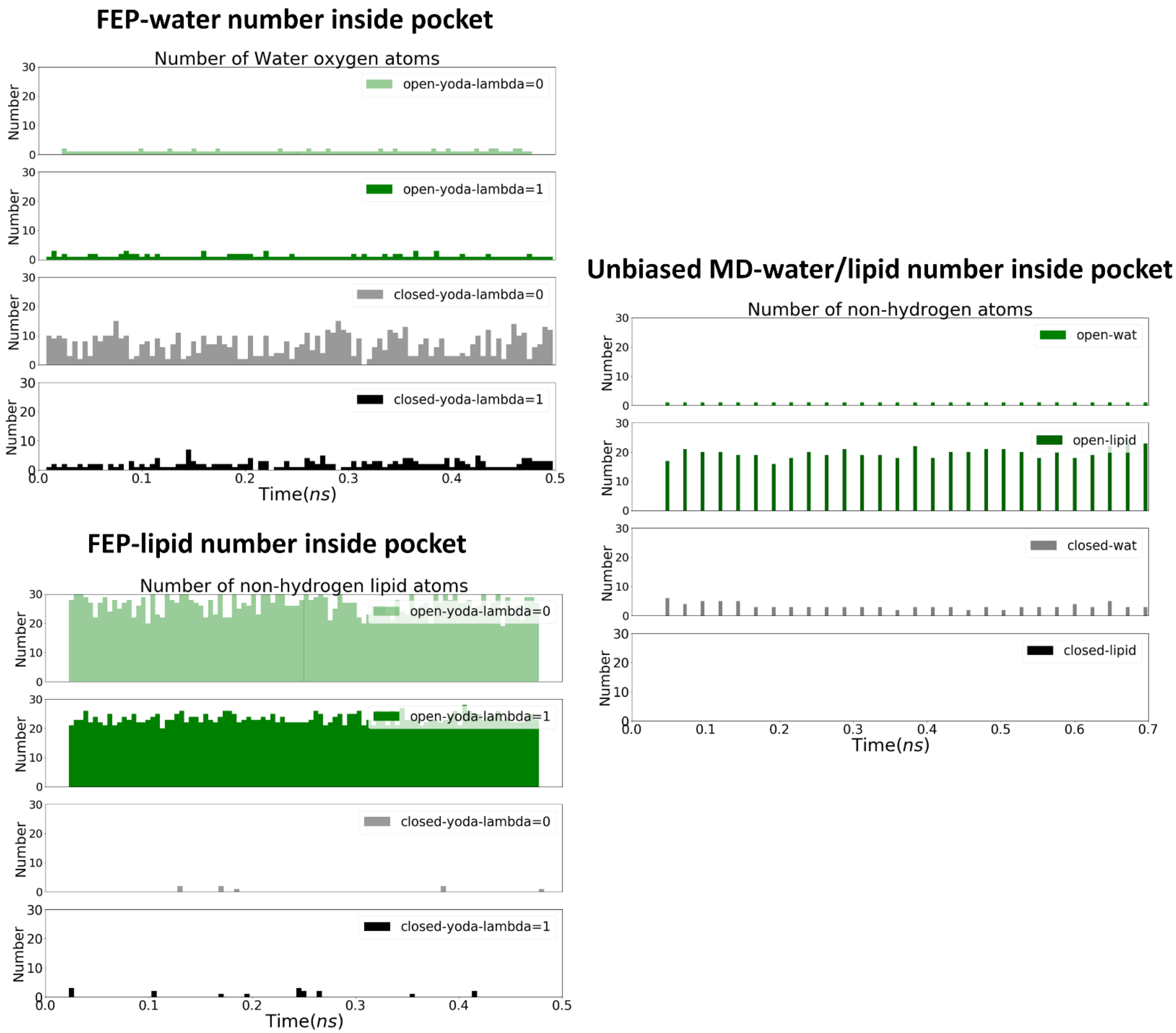


**Figure S9**. The time series of the number of water oxygen and non-hydrogen lipid atom inside the Yoda1 binding pocket during FEP/REMD simulations and unbiased simulations without Yoda1. The pocket is defined by a cutoff distance of 8Å from the center of mass of the binding pocket (residue ID: 1683-1733, 2047-2097).

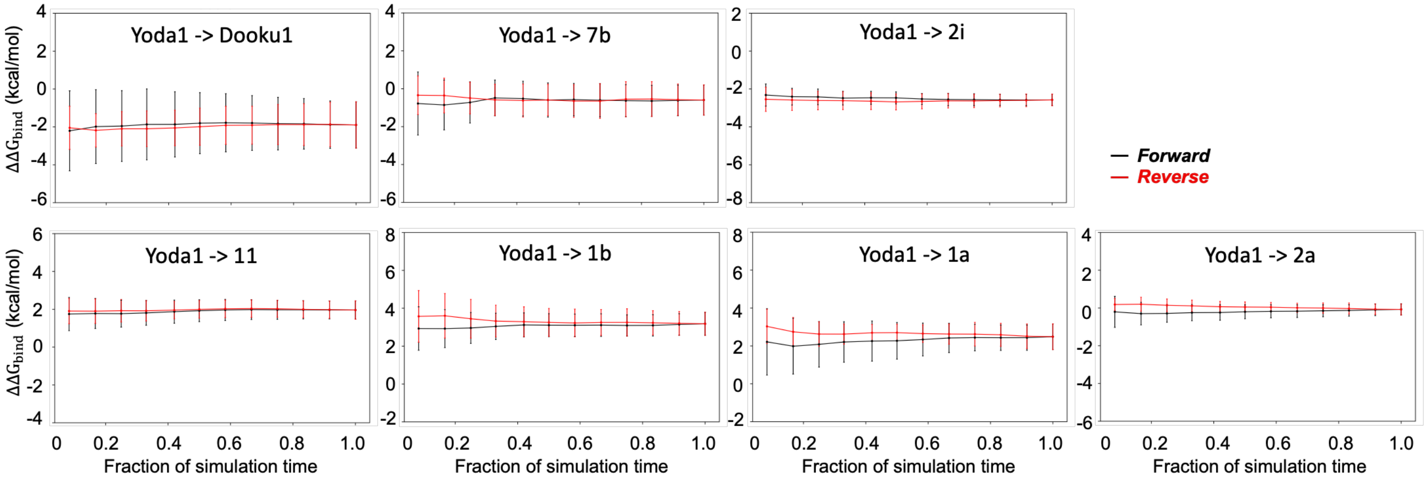


**Figure S10.** Free energy convergence as a function of time for the seven transformation pairs with a 12 λ window scheme (0.0000, 0.0479, 0.1151, 0.2063, 0.3161, 0.4374, 0.5626, 0.6839, 0.7937, 0.8850, 0.9521, and 1.0000) in the open state. 5 ns simulations were performed in each window. The $\Delta\Delta G_{\mathrm{bind}}$ convergence is estimated from the froward (black) and time-reversed (red) data of the last 4 ns trajectory.

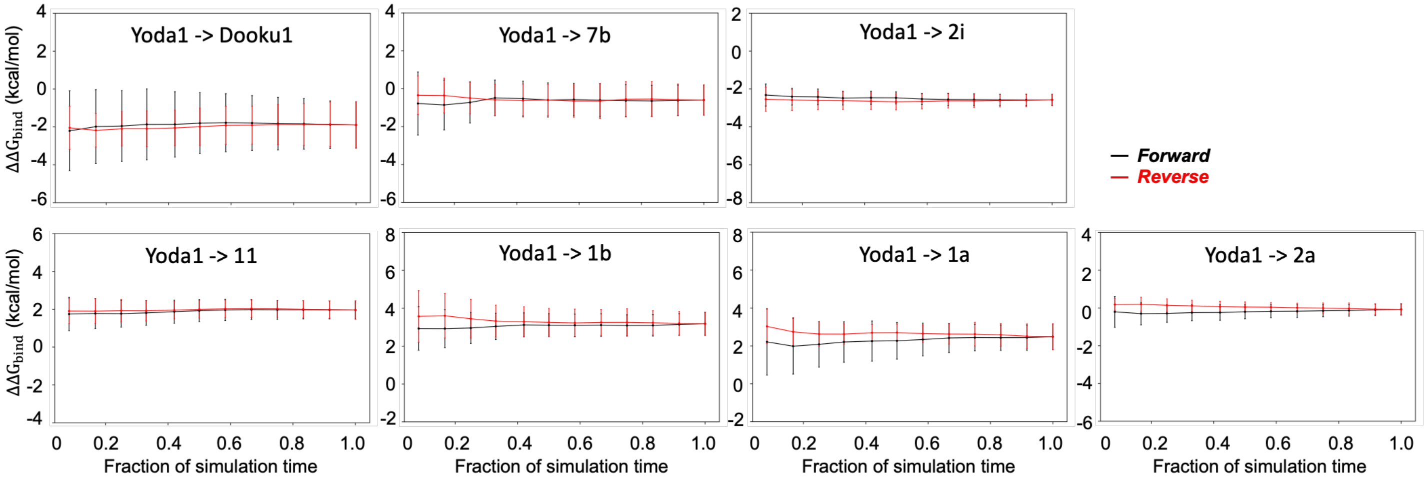


**Figure S11.** Free energy convergence as a function of time for the seven transformation pairs with a 12 λ window scheme (0.0000, 0.0479, 0.1151, 0.2063, 0.3161, 0.4374, 0.5626, 0.6839, 0.7937, 0.8850, 0.9521, and 1.0000) in the closed state. 5 ns simulations were performed in each window. The $\Delta\Delta G_{\mathrm{bind}}$ convergence is estimated from the froward (black) and time-reversed (red) data of the last 4 ns trajectory. To ensure the convergence of our AMBER-TI simulations, we extend 10 ns simulations per window for the transformation pair, Yoda1 -> Dooku1. The $\Delta\Delta G_{\mathrm{bind}}$ convergence is estimated from the froward and time-reversed data of the last 5 ns trajectory for Yoda1 -> Dooku1.
